# Green algae CO_2_ capture is powered by alternative electron pathways of photosynthesis

**DOI:** 10.1101/2023.08.08.552514

**Authors:** Gilles Peltier, Carolyne Stoffel, Justin Findinier, Sai Kiran Madireddi, Ousmane Dao, Virginie Epting, Amélie Morin, Arthur Grossman, Yonghua Li-Beisson, Adrien Burlacot

## Abstract

On Earth, microalgae contribute to about half of global net photosynthesis. During photosynthesis, sunlight is converted into chemical energy (ATP and NADPH) used by metabolism to convert CO_2_ into biomass. Alternative electron pathways of photosynthesis have been proposed to generate additional ATP that is required for sustaining CO_2_ fixation, but the relative importance of each pathway remains elusive. Here, we dissect and quantify the contribution of cyclic, pseudo-cyclic and chloroplast to mitochondria electron flows for their ability to sustain net photosynthesis in the microalga *Chlamydomonas reinhardtii*. We show that each pathway has the potential to energize substantial CO_2_ fixation, can compensate each other, and that the additional energy requirement to fix CO_2_ is more than 3 times higher than previous estimations. We further show that all pathways have very different efficiencies at energizing CO_2_ fixation, with the chloroplast-mitochondria interaction being the most efficient, thus laying bioenergetic foundations for biotechnological improvement of CO_2_ capture.

## Introduction

Oxygenic photosynthesis is the main process responsible for the carbon input into ecosystems on Earth, with net CO_2_ fixation by photosynthesis representing more than ten times anthropic CO_2_ emissions (IPCC, 2021). During photosynthesis, carbon fixation is powered by the combined action of photosystems (PS) I and II, establishing a linear electron flow (LEF) that produces reducing power (NADPH) and a trans-thylakoidal proton motive force (*pmf*) used to generate ATP. Both ATP and NADPH fuel the CO_2_ fixation reactions of the Calvin-Benson-Bassham (CBB) cycle and the further processing of the glyceraldehyde-3-phosphate (G3P) to regenerated ribulose-1-5 bisphosphate and produce building blocks used for the elaboration of biomass (Johnson and Alric, 2013). However, given the insufficiency of the LEF to generate an adequate *pmf* and the conversion efficiency of the chloroplastic ATP-synthase (ATPase), it has been long recognized that the LEF only generates 85% of the theoretical ATP requirements of the CBB cycle (Allen, 2003).

Cyclic electron flow, which recycles reducing equivalents around PSI (CEF) and pseudo-cyclic electron flow (PCEF), which reduces molecular O_2_ at the acceptor side of PSI (**Fig. 1**), were proposed to contribute to the production of extra ATP during photosynthesis (Arnon, 1959, 1984). In most angiosperms and mosses, CEF entails two main pathways, one involving the proton gradient generation (PGR) 5 (Munekage *et al*., 2002) and PGR-Like 1 proteins (DalCorso *et al*., 2008) and another involving the plastidial NDH-1 complex (Joet *et al*., 2001; Munekage *et al*., 2004). In green microalgae, the plastidial NDH-1 complex is absent (Peltier *et al*., 2016) and it was reported that two CEF pathways operate, one controlled by PGRL1 and PGR5 (Tolleter *et al*., 2011; Johnson *et al*., 2014) and the other involving the plastidial NAD(P)H dehydrogenase Nda2 (Jans *et al*., 2008; Desplats *et al*., 2009; Saroussi *et al*., 2016). Since the Nda2 pathway is mostly active during nitrogen deprivation (Peltier and Schmidt, 1991; Saroussi *et al*., 2016), hereafter CEF refers to the PGRL1/PGR5-controled CEF. PCEF involves Flavodiiron proteins (FLVs) that catalyse O_2_ photoreduction at the PSI acceptor side (**Fig. 1**) in cyanobacteria(Helman *et al*., 2003; Allahverdiyeva *et al*., 2013), green microalgae (Chaux *et al*., 2017), mosses (Gerotto *et al*., 2016), liverwort (Shimakawa *et al*., 2017) and gymnosperms (Ilík *et al*., 2017).

**Figure 1.**
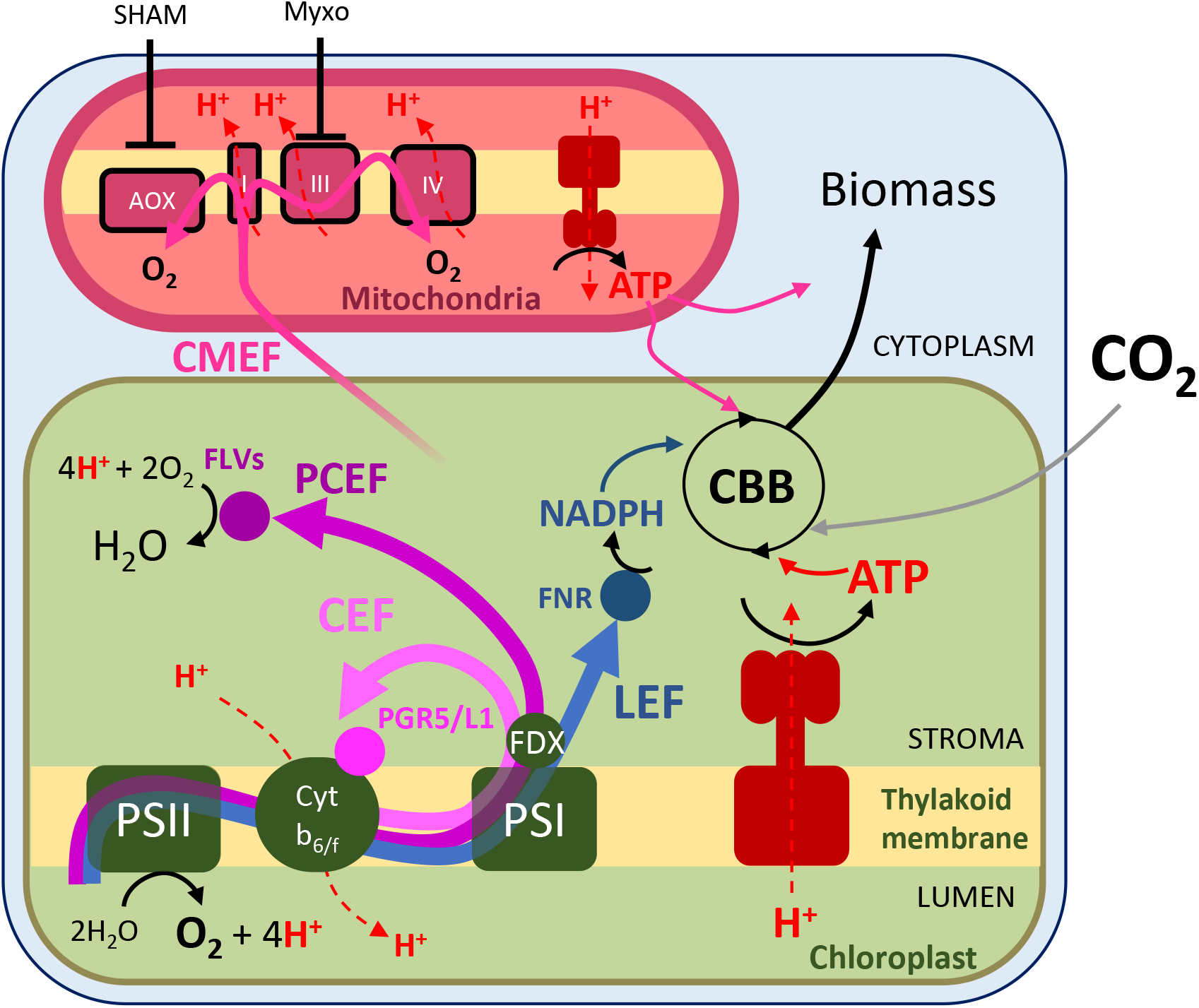
Electron transport pathways operating during oxygenic photosynthesis and their role in ATP generation. Linear electron flow (LEF) operates from PSII to PSI and reduces NADP^+^ to NADPH using electrons from water. It also generates a *pmf* across the thylakoid membrane composed of a proton gradient (ΔpH) and electrical gradient (ΔΨ); the *pmf* is used by the ATPase to generate ATP. Cyclic electron flow (CEF) and pseudo cyclic electron flow (PCEF) both contribute to the generation of the *pmf* but do not produce reducing equivalents since the e^-^ used for PCEF reduce O_2_ to water. Chloroplast to mitochondria electron flow (CMEF) encompasses multiple metabolic pathways by which reductants generated in the chloroplast is shuttled to the mitochondria and converted into ATP by the mitochondrial electron transport chain. NADPH and ATP generated by these different pathways are eventually used for transforming CO_2_ into biomass, which reflects net photosynthesis. Other abbreviations: AOX, alternative oxidase; ATP, adenosine triphosphate; CBB, Calvin Benson Bassham; FDX, ferredoxin; FLVs, flavodiiron proteins; FNR, ferredoxin-NADP^+^ reductase; Myxo, myxothiazol; NADPH, reduced form of nicotinamide adenine dinucleotide phosphate; PS, photosystem; PGR5/L1, proton gradient generation 5/like-1; SHAM, salicyl hydroxamic acid.

Another mechanism that can supply ATP to photosynthesis involves metabolic cooperation between chloroplasts and mitochondria (Raghavendra and Padmasree, 2003). Reducing power generated within the chloroplast is transferred to the mitochondria by metabolic shuttles where it is converted into ATP by the mitochondrial respiratory chain and might be shuttled back to the cytosol and the chloroplast (**Fig. 1**). Such a cooperation, called here Chloroplast to Mitochondria Electron Flow (CMEF) was first evidenced from the restoration of photoautotrophic growth in a mutant of the green microalgae *Chlamydomonas reinhardtii* (hereafter *Chlamydomonas*) deficient in the plastidial ATPase (Lemaire *et al*., 1988). Similar cooperation was later suggested from the study of *Chlamydomonas* mutants affected in mitochondrial respiration (Cardol *et al*., 2003) or in CEF (Dang *et al*., 2014) as well as in the moss *Physcomytrum patens* (Mellon *et al*., 2021), and in diatoms (Bailleul *et al*., 2015).

It is generally considered that both PCEF and CMEF act essentially as electron valves, operating either transiently in the case of PCEF to alleviate excess electrons generated by photosynthesis under light fluctuating regimes (Alric and Johnson, 2017; Chaux *et al*., 2017; Alboresi *et al*., 2019) or more continuously for CMEF under high light stress (Kaye *et al*., 2019). Although in *Chlamydomonas*, CMEF, PCEF and CEF have been recently shown to all contribute to supply energy to the CO_2_ concentrating mechanism (CCM) (Burlacot *et al*., 2022), evidence of their role in powering the CBB cycle and downstream carbon metabolism processes are scarce.

In this work, we show that each pathway (CEF, PCEF and CMEF) has the capacity to sustain net CO_2_ fixation to varying degrees and can compensate for the absence of the other pathways, whereas CO_2_ fixation is completely abolished in the absence of all three pathways. Moreover, we introduce a new metric: the electron quantum yield of net photosynthesis for each pathway and use it to show that CMEF is by far the most efficient at energizing CO_2_ fixation. We further discuss our data in relation to the relative contribution and efficiency of the different alternative pathways to the net primary productivity of terrestrial and oceanic ecosystems.

## Results

### Net photosynthesis fully relies on mitochondrial respiration when PCEF and CEF are impaired

To test whether PCEF and CEF are required for maximal net photosynthesis (V_Max_), we first measured photosynthetic O_2_ exchange rates using Membrane Inlet Mass Spectrometry (Burlacot *et al*., 2020) in air-grown *Chlamydomonas* mutants impaired in both CEF and PCEF (*pgrl1 flvB-1 and 3*) and in control strains (WT2 and 3)(Burlacot *et al*., 2022). Since the *pgrl1* and *flvB* mutants are affected in the functioning of the CO_2_ concentration mechanism (CCM) (Burlacot *et al*., 2022), photosynthesis measurements were performed in the presence of a saturating concentration of inorganic carbon (C_i_) when the CCM is inactive. As previously reported (Burlacot *et al*., 2022), *pgrl1 flvB* double mutants can perform photosynthesis at maximum rates comparable to the single *pgrl1* or *flvB* mutants and to the parental and sibling control strains (**Fig. 2; Sup Figs. S1 A, S2, S3**), therefore showing that neither CEF nor PCEF are essential to reach V_Max_. To test a possible contribution of mitochondrial respiration to photosynthesis, *pgrl1 flvB* double mutants were treated with myxothiazol (Myxo) and salicylhydroxamic acid (SHAM), inhibitors of mitochondrial electron transport, respectively acting on complex III and the alternative oxidases (AOX) (**Fig. 1**)(Cournac *et al*., 2002). While the addition of both inhibitors had minimal impact (about 10-20% decrease) on the V_Max_ of control strains (**Fig. 2 A, C, Sup Figs. S1 A, S2, S3**), they completely abolished photosynthesis of *pgrl1 flvB* double mutants (**Fig. 2 B, C, Sup Figs. S1, S2)**.

**Figure 2.**
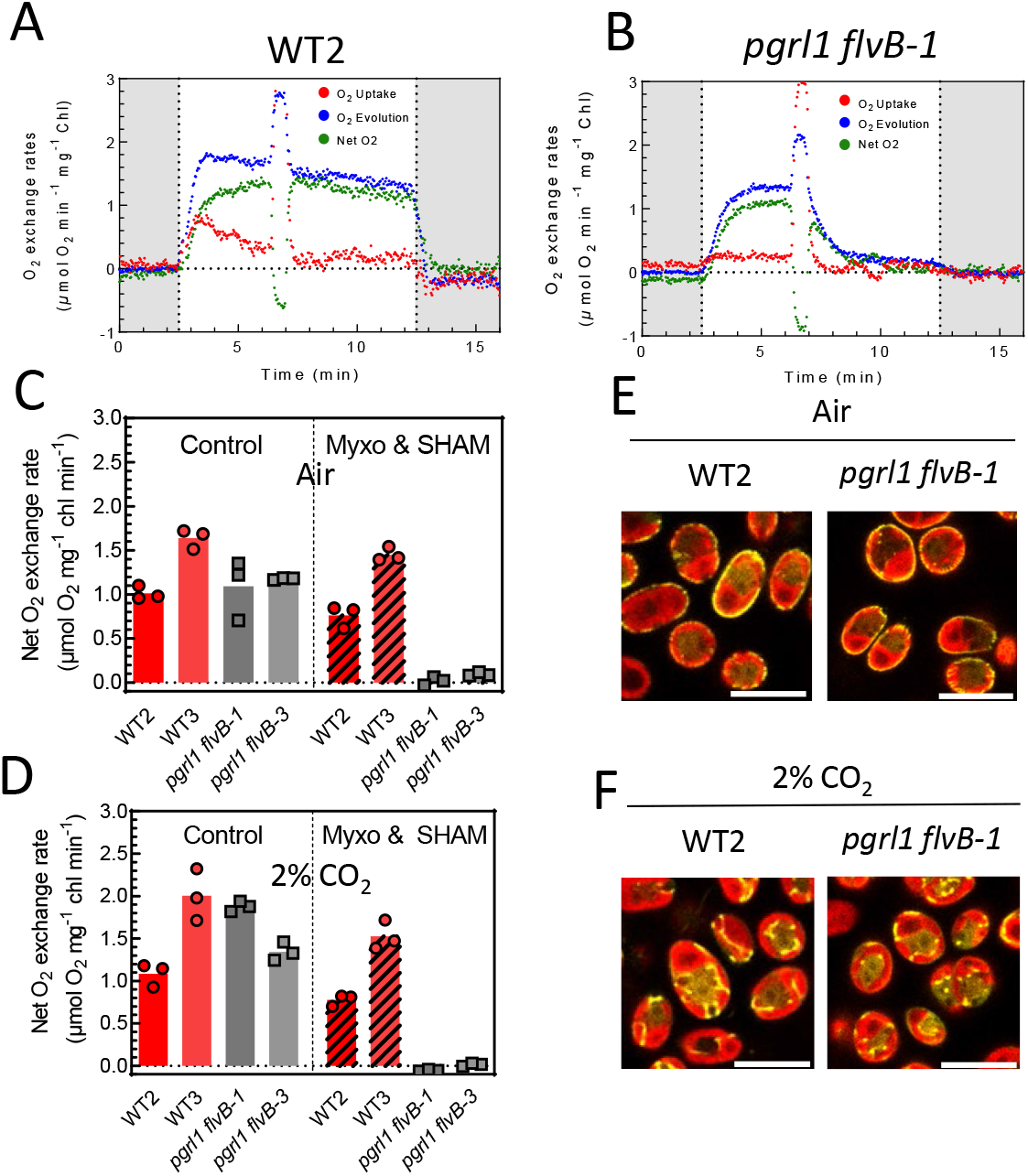
Dependence of net photosynthesis on CEF, PCEF and CMEF. **A, B**, O_2_ exchange rates upon a dark to light transition in WT2 (**A**) and *pgrl1 flvB-1* (**B**). At t=6min, mitochondrial respiration inhibitors myxothiazol (2.5L*µ*M) and salicylhydroxamic acid (SHAM, 400L*µ*M) were added to the reaction. Shown are representative traces of *n*L=L3 biologically independent samples. **C and D**, Net O_2_ evolution in *pgrl1 flvB-1* and *pgrl1 flvB-*3 and their control strains (WT2 and WT3) in the absence (control) or presence (Myxo & SHAM) of myxothiazol (2.5L*µ*M) and salicylhydroxamic acid (SHAM, 400L*µ*M) in cells grown in air (**C**) or in 2% CO_2_-enriched air (**D**). Values are taken after 9 min of illumination upon a dark to light transitions. Bars show the average and dots show individual replicates (*n*L=L3 biologically independent samples). Typical experiments used to extract these data are shown in **Sup. Fig. S2**. **E** and **F**, Cross section images showing subcellular localization of mitochondria-localized Clover (yellow fluorescence) in *pgrl1 flvB-1* and a control strain (WT2), grown in air (**E**) or 2% CO_2_-enriched air (**F**). Images are representative of 2 independent experiments. Scale bar: 10 µm. Brightfield, Clover fluorescence, Chlorophyll autofluorescence and Clover/Chlorophyll merge signals are shown. Images are representative of 2 experiments. Scale bar: 10 µm.

In *Chlamydomonas*, the cellular location of mitochondria depends on the CO_2_ supply, with the mitochondria located at the cell periphery in low CO_2_-grown cells and mostly inside the chloroplast cup in high CO_2_ (Geraghty and Spalding, 1996) (**Fig. 2E, F**). Since this location might affect CMEF (Burlacot and Peltier, 2023), we sought to examine the involvement of the mitochondrial sub-cellular localization on the role of CMEF. Similar to air-grown cells, mitochondrial inhibitors fully stopped net photosynthesis on 2% CO_2_-grown *pgrl1 flvB* mutants while having a marginal effect (about 10-20% decrease) on the control strains (**Fig. 2D, Sup Fig. S4**). To control the location of the mitochondria in the different strains, we visualized the spatial distribution of an expressed, mitochondria-targeted fluorophore (Clover). There was similar re-localization of mitochondria depending on the CO_2_ supply during growth (at periphery under very low CO_2_) of both the single and double mutants and a control strain (**Fig. 2E, F, Sup Fig. S5**). We conclude from this experiment that in the absence of both CEF and PCEF, near wild type levels of photosynthesis can be achieved that fully rely on CMEF, the role of CMEF being independent of the spatial distribution of the mitochondria.

### CEF, PCEF and CMEF each have the potential to sustain most net photosynthesis

We quantified the potential of CEF, PCEF and CMEF to energize net photosynthesis, by measuring the effect of mitochondrial inhibitors on photosynthesis rates in single and double *pgrl1* and *flvB* mutants (**Fig. 3 A, B, C, D**). Inhibition of respiration decreased net photosynthesis by 40-50% in both the *flvB* and *pgrl1* single mutants (**Fig. 3 A, B, Sup Figs. S1 A, S2**), showing that both CEF and PCEF have the potential to energize 50-60% of V_Max_ on their own (**Fig. 3 E, Sup Fig. S1A, B**). Inhibition of respiration decreased the V_Max_ of all control strains by 10-20% (**Fig. S1 A, S2, S3**) showing that combined activities of CEF and PCEF can power approximately 85% of photosynthesis (**Fig. 3 E**). Based on the effects of SHAM or myxothiazol individually, on V_Max_ in *pgrl1 flvB* mutants (**Fig. 3C, D, E, Sup Figs. S1 A, S2**), we conclude that while the complex III-IV pathway of CMEF (**Fig. 1**) can power most net photosynthesis on its own (**Fig. 3D**), the AOX pathway (**Fig. 1**) can only sustain approximately 45% of V_Max_ (**Fig. 3E**). Because the amount of the terminal oxidases involved in CMEF does not differ in the *pgrl1 flvB* mutants and their control strains (**Sup Fig. S6**), these values likely reflect wild type capacities. We also quantified net photosynthesis based on inorganic carbon (C_i_) uptake (**Sup Figs. S7, S8**). On all strains tested, the inhibitory effects on V_max_ measured as net C_i_ uptake was similar to that observed on net O_2_ production (**Sup Figs. S7**). We conclude from these experiments that CEF, PCEF and CMEF can sustain photosynthetic C_i_ uptake on their own but that only the complex III-IV pathway of CMEF is efficient enough to reach near maximal photosynthesis levels.

**Figure 3.**
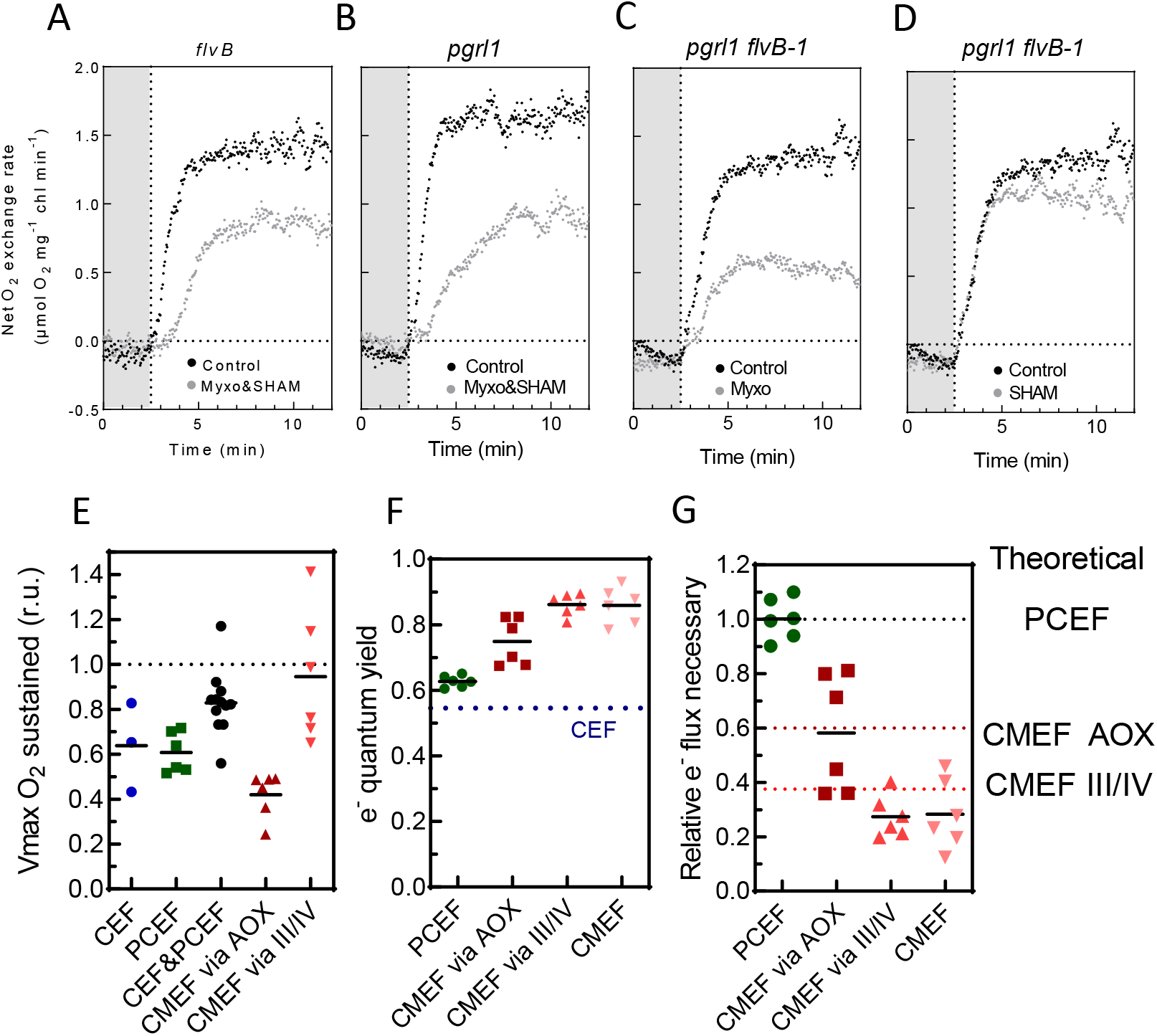
Relative efficiencies of CEF, PCEF and CMEF in energizing net photosynthesis. **A-D**, Net O_2_ evolution measured during a dark to light transition in *flvB* (**A**), *pgrl1* (B), *pgrl1 flvB*-2 (**C**, **D**), treated (gray dots) or not treated (black dots) with both myxothiazol (Myxo, 2.5L*µ*M) and salicylhydroxamic acid (SHAM, 400L*µ*M) (**A, B**), or with only one inhibitor (**C,D**). Shown are representative traces of 3 biological replicates. **E**, Relative V_max_ measured when a given set of alternative electron pathways are active after treatment with myxothiazol (Myxo, 2.5L*µ*M), salicylhydroxamic acid (SHAM, 400L*µ*M) or both inhibitors; CEF (blue, *flvB* treated with Myxo and SHAM), PCEF (green, *pgrl1* treated with Myxo and SHAM), CEF&PCEF (dark, control strains treated with Myxo and SHAM), CMEF via AOX (purple, *pgrl1 flvB* treated with Myxo), CMEF via III/IV (red, *pgrl1 flvB* treated with SHAM). **F,** Electron quantum yield of PCEF (green), CMEF via AOX (purple), CMEF via III/IV (red), CMEF (pink) and CEF (dotted line). Similar electron quantum yields were also determined for PCEF in 2% CO_2_-grown cells and a slightly lower quantum yield for CMEF (**Sup Fig. S9 D**). **G**, Electron flux relative to PCEF required to power similar levels of net photosynthesis when relying on a single pathway. Data shown are individual extrapolations based on measured electron quantum yields. In **E, F** and **G,** bars show the average and dots show individual replicates (*n*L≥3 biologically independent sample).

### CEF, PCEF and CMEF show different efficiencies in sustaining net photosynthesis

Mechanistically, for each electron flowing once through the different pathways (CEF, PCEF, the AOX, or the complex III-IV pathway of CMEF), the total number of protons translocated across the thylakoid and/or the inner mitochondrial membranes, highly differ (Burlacot, 2023). To quantify the potential of the different pathways in energizing net photosynthesis, we introduce a new metric, the electron (e^-^) quantum yield of a pathway defined by the following equation:

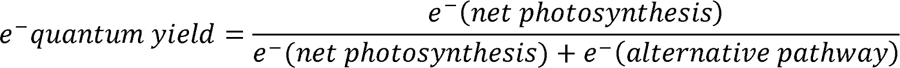

Where e^-^ (net photosynthesis) is the flux of photosynthetic electrons directly used for net photosynthesis and e^-^ (alternative pathway) is the flux of photosynthetic electrons through a given alternative pathway under conditions where only this alternative pathway is active. Therefore, the e-quantum yield of an alternative pathway, which is determined when only this alternative pathway is operating, characterizes the efficiency of electrons passing through it in powering net photosynthesis. Note that the e^-^ quantum yield, thus does not reflect the rate or the importance of the pathway considered but rather its capacity to produce ATP for each electron. We used this equation and experiments from **Fig. 3 A-E** to quantify the e^-^ quantum yield of each pathway (**Fig. 3 F, Sup Fig. S9**). The e^-^quantum yield of PCEF was estimated at 0.65, whereas that of CMEF at 0.85 (**Fig. 3F**), indicating a better efficiency of CMEF in driving CO_2_ fixation as compared to PCEF. In the absence of an established method to quantifying CEF in physiologically relevant conditions (Burlacot *et al*., 2022), CEF quantum efficiency was estimated to be around 0.55 by hypothesizing that an electron passing once through CEF is 1.5 time less efficient at generating a *pmf* than if it would pass through PCEF (Kramer and Evans, 2011; Burlacot, 2023) (**see Materials and Methods**). In the case of CMEF, the e^-^ quantum yield of the AOX pathway was lower than that of the complex III-IV pathway (**Fig. 3F**), likely reflecting the proton translocating capacity of each pathway (**Fig. 1)(Burlacot, 2023)**. Using the e^-^ quantum yields of each pathway, we then compared the flux of electrons required to power the same amount of net photosynthesis relative to PCEF (**Fig. 3G**). Strikingly, theoretical values were in accord with values experimentally determined by the e^-^ quantum yields (**Fig. 3 G**). We conclude from these experiments that CMEF pathways are more efficient than PCEF and CEF at energizing CO_2_ fixation and that this difference arises from a higher H^+^/e^-^ transfer capacity likely leading to a higher capacity to generate ATP.

### Estimating the respective contributions of CEF, PCEF and CMEF during steady state photosynthesis and growth

We then sought to estimate (*i*) the electron flux via each pathway expressed as an additional electron flow relative to net photosynthesis (*%-LEF*) and (*ii*) their contribution to energize net photosynthesis. We used two different approaches, one based on the effect of mutations, the other based on the effect of respiratory inhibitors. As both approaches are based on pathway inhibition, they are potentially biased by the high level of redundancy existing between the different AEFs and the existence of compensatory mechanisms (Cardol *et al*., 2009; Dang *et al*., 2014; Burlacot *et al*., 2022). The two approaches differ, however, in the way compensatory mechanisms likely contribute, mutants being more subject to compensation than short-terms effects of inhibitors. In the mutant-based approach, PCEF was quantified by subtracting O_2_ uptake rates in the *flvB* mutant from its control parental strain (allowing the calculation of PCEF_mut_ and CMEF_mut_) (**Sup Fig. S10**). In the inhibitor-based approach, CMEF was quantified by subtracting O_2_ uptake rates between SHAM plus Myxo treated and untreated control strains (allowing the calculation of PCEF_inhib_ and CMEF_inhib_) (**Sup Fig. S10**). The two methods differ somewhat in their estimates of PCEF and CMEF (**Fig. 4 A, B, Sup Fig. S11**), with a higher CMEF contribution being estimated using the mutant method. On average, PCEF and CMEF represent a flux of between 10-20*%-LEF* and 5-10*%-LEF*, respectively (**Fig. 4 A**). Thanks to the respective e^-^ quantum yield of each pathway, we can estimate that 10-30% and 30-60% of CO_2_ fixation is sustained by PCEF and CMEF respectively (**Fig. 4 B**). Since CEF sustains the remaining net photosynthesis, we estimated that 25-30% of CO_2_ fixation is powered by CEF (**Fig. 4 B**), which corresponds to a CEF of about 25*%-LEF* assuming an e^-^ quantum yield of CEF of 0.55 (**Fig. 4 A**).

**Figure 4.**
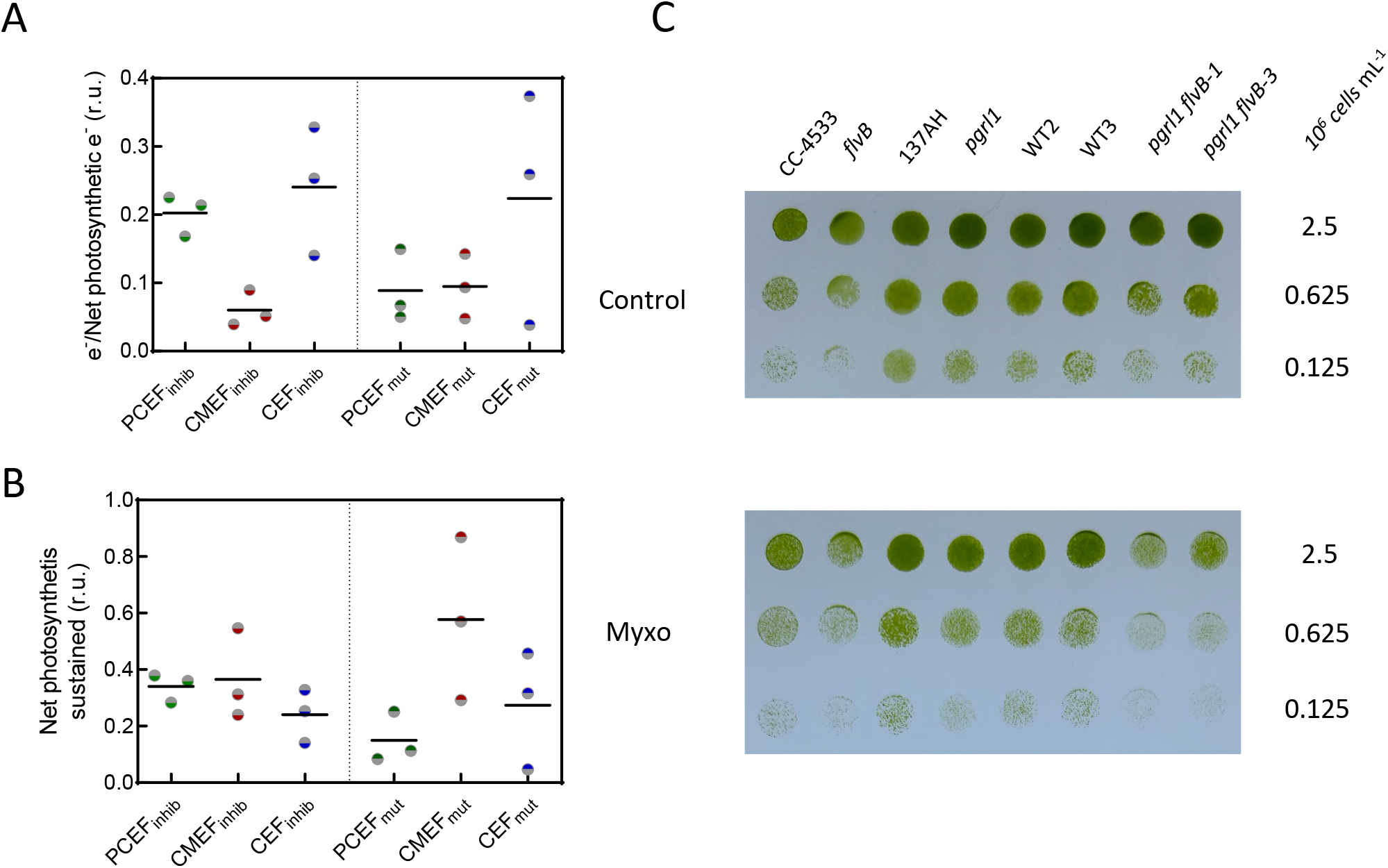
Contribution of CEF, PCEF and CMEF to the electron flow, net photosynthesis and growth. **A** Electron flow through PCEF, CEF and CMEF relative to LEF were determined in the wildtype strain CC-4533 by two different methods, one based on the use of inhibitors (inhib) the other on mutants (mut). Data used for these calculations are the same as for Fig. 2 C and **Sup Fig. 2. B** Proportion of net photosynthesis powered by each electron flow (PCEF, CEF and CMEF) in the control strain CC-4533 as measured by the inhibition (inhib) or mutant (mut) method. CC-4533 was chosen because it is the control strain for *flvB* and is therefore the only strain where both methods could be used. Data used for these calculations are the same as for **A**. In **A** and **B,** bars show the average and dots show individual replicates (*n=*3 biologically independent sample). **C**, Growth tests for *flvB* and *pgrl1* mutants, their corresponding control strains (CC-4533 and 137AH, respectively) and double mutants (*pgrl1 flvB-1 and pgrl1 flv-3*) and their corresponding control strains (WT2 and 3). Cells were spotted on plates containing minimal medium at pH 7.2 and grown under continuous illumination (60Lµmol photon m^−2^ s^−1^) in the absence (upper panel) or presence (lower panel) of 2.5µM Myxothiazol (Myxo) in air enriched at 2% CO_2_. Spots shown are representative of 10 independent experiments.

To further assess the contribution of CEF, PCEF and CMEF over the long term, growth tests were performed on the different strains in the absence or presence of the respiration inhibitor Myxo. While Myxo has no effect on the growth of control strains and single *flvB* or *pgrl1* mutants in moderate light intensities (**Fig. 4 C**), growth of *pgrl1 flvB* was impaired in the presence of the inhibitor (**Fig. 4 C**). Interestingly, at low light intensity (30 µmol photons m^-2^ s^-1^), the effect of Myxo on the double mutant was absent (**Sup Fig. 12 A**), suggesting that the AOX pathway of CMEF has the potential to fully power net photosynthesis in low light conditions. Therefore, whereas CEF, PCEF, and CMEF all contribute to energizing photosynthesis, CMEF, although representing a relatively small flux, has the potential to fuel CO_2_ fixation and biomass accumulation thanks to its high e^-^ quantum yield.

## Discussion

Since the first description of CEF by Arnon in 1959 (Arnon, 1959), the question was raised as to whether LEF could sustain the ATP requirement of photosynthetic CO_2_ fixation (Arnon, 1959), and it was proposed that photosynthetic electron flows, other than LEF, are involved in balancing the ATP/NADPH ratio to sustain CCB cycle function (Arnon, 1959; Allen, 2003). Since then, several mechanisms of CEF, PCEF and CMEF have been uncovered (Munekage *et al*., 2002; Helman *et al*., 2003; Munekage *et al*., 2004; Tomaz *et al*., 2010) as well as some compensations between those different mechanisms (Cardol *et al*., 2009; Petroutsos *et al*., 2009; Dang *et al*., 2014; Storti *et al*., 2019; Storti *et al*., 2020; Burlacot *et al*., 2022), but the importance and relative efficiency of each mechanism remain mostly unknown. Here we have shown that in *Chlamydomonas* PGRL1-controlled CEF, FLV-mediated PCEF, and both the AOX and complex III/IV/cytochrome oxidase pathways of CMEF can all supply additional energy to complement LEF for CO_2_ fixation. The e^-^ quantum yields of CEF, PCEF and CMEF greatly differ; CMEF is the most electron efficient and contributes importantly to the extra ATP requirements of CO_2_ fixation in continuous, saturating light. Note, that in spite of the ongoing controversy on how PGRL1 is involved in CEF (Nawrocki *et al*., 2019), a clear integration is observed between PGRL1-controled CEF, PCEF and CMEF in which these different pathways can compensate each other in powering photosynthesis. Importantly, we observed similar effects on net O_2_ and CO_2_ exchange (**Sup Fig. S7**), thus showing that all effects measured on net photosynthesis correspond to changes in CO_2_ fixation and eventually biomass production and growth (**Fig. 4 C**).

Some compensation between CEF, PCEF, and CMEF has been previously reported; a *Chlamydomonas* PGRL1-deficient strain exhibits increased CMEF and PCEF (Dang *et al*., 2014) and mutants of complexes I and III show enhanced rates of CEF (Cardol *et al*., 2009). We observed an elevated CMEF in the absence of FLV (**Fig. 4 A, B**) with a higher level of terminal oxidases associated with CMEF (**Sup Fig. 6**), corroborating cross-compensation among all three mechanisms of electron flow. In diatoms, while CMEF is considered to be a major driver of CO_2_ fixation, inhibition of CMEF only leads to a ∼20% decrease of the PSII yield (Bailleul *et al*., 2015), which might also reflect compensation by CEF. We also conclude, based on the fact that no CO_2_ fixation could be detected in the double *pgrl1 flv* mutants treated with inhibitors of mitochondrial respiration, that neither the NDA2-mediated CEF, nor O_2_ reduction mediated by the plastid terminal oxidase or Mehler reactions have the potential by themselves to sustain a measurable amount of net photosynthesis, which is in agreement with previous studies concluding that these pathway represent a small fraction of alternative electron flow (Alric *et al*., 2010; Houille-Vernes *et al*., 2011; Nawrocki *et al*., 2019)(**Fig. 2 B**).

The quantitative contribution of each pathway in the wild-type strain in the absence of perturbation of any AEFs is a challenging task to which the approaches we have developed here provide an imperfect answer. Indeed, the quantification of each pathway we have proposed relies on the assumption that inhibiting one pathway (CMEF or PCEF) does not affect the flow through the others, assumption which is most likely not verified since compensation between different AEFs mechanisms have been previously documented (Cardol *et al*., 2009; Dang *et al*., 2014; Burlacot *et al*., 2022). It should therefore be considered as a first attempt to evaluate the respective contributions of each pathway until an integrative quantification method is available. Here, the mutant method likely overestimates CMEF and the inhibitor method likely overestimates PCEF, both approaches likely overestimating CEF. It is clear however, that no single pathway sustains net photosynthesis by itself (**Sup Fig. S11 B**) which might reflect a rather balanced contribution of the different pathways to CO_2_ fixation in the WT. In any case, it seems highly likely that, considering the high redundancy of AEFs and the ability of each of them to sustain CO_2_ fixation at a high rate, a dynamic mix of the different pathways may be involved in different environmental conditions, each pathway being more or less advantageous in specific conditions (Burlacot, 2023).

### The e^-^ quantum yield, a new tool to evaluate the efficiency of electron transport pathways

In support to LEF, a combination of PGRL1-controled CEF and PCEF cannot sustain more than 80% of maximal CO_2_ fixation (**Fig. 3 E**) whereas each individual flux can help sustain 60% of it (**Fig. 3 F**). This suggests the existence of an intrinsic limitation of the flow of electrons through the photosynthetic electron transport chain, which is likely due to a *pmf*-induced photosynthetic control occurring at the cytochrome *b_6_f* level (Johnson and Berry, 2021; Malone *et al*., 2021); an elevated *pmf* eliciting a block in electron flow across the *b_6_f* complex. Engaging an electron pathway with a high e^-^ quantum yield, such as CMEF, might be a way to avoid this limitation since less electrons would be needed to power the same level of CO_2_ fixation. Such a strategy seems to be used by diatoms (Bailleul *et al*., 2015) which lacks FLVs.

Throughout the green lineage, the nature and activity of alternative electron flows are quite different. For example, angiosperms lack FLV-dependent PCEF (Peltier *et al*., 2010), harbour a NDH-1-dependent CEF pathway (Peltier *et al*., 2016) and have a weakly active CMEF (Raghavendra and Padmasree, 2003). Since the NDH-1 dependent CEF translocates 4H^+^/e^-^ (Strand *et al*., 2017), its e^-^ quantum yield is close to that of CMEF, strongly suggesting that in angiosperms the NDH-1 complex can compensate when the CMEF capacity is diminished. The latter is illustrated by the poor growth of *Arabidopsis thaliana* mutants impaired in both CEF pathways (PGR and NDH-1) (Munekage *et al*., 2004). Because of the number of key proteins and biochemical reactions involved in CMEF, its high e^-^ quantum yield may not be as rapidly accessible as CEF or PCEF during environmental fluctuations. While under light limiting conditions a high e^-^ quantum yield pathway might be advantageous, a less efficient pathway would contribute to the safe dissipation of excess absorbed light energy and diminished reactive oxygen production at saturating light intensities. Overall, it remains to be determined to what extent harbouring a palette of alternative electron flows with varying quantum yields and likely different time responses, is critical for sustaining photosynthetic performance under dynamic conditions in nature.

Note here that while modulation of the capacity of membranes to transform *pmf* into ATP may modulate the absolute e^-^ quantum yield of each pathway, the relative e^-^ quantum yield of CEF and PCEF would not change if they are embedded in the same thylakoid system. Similarly, even in conditions where the mitochondrial respiration would be partially or fully uncoupled, the e^-^ quantum yield of CMEF would be at least as good as PCEF due to the generation of a *pmf* across the thylakoid membranes (Burlacot, 2023). Hence, if there is no electron transport chain dedicated to a single pathway, electrons passing through CMEF and PCEF should always be more efficient than CEF in powering net photosynthesis.

### Additional energy cost of photosynthesis

Allen previously estimated that a PCEF representing only 16*%-LEF* would be sufficient, in addition to LEF, to satisfy the ATP and NADPH requirement of the CBB cycle (Allen, 2002). Our conclusions are strikingly different since we estimate that if only PCEF was active, it would have to represent 54*%-LEF* to sustain CO_2_ fixation. This discrepancy may reflect the fact that the actual metabolic cost of CO_2_ fixation exceeds the ATP requirement of the CBB cycle. This may result from the involvement of additional ATP-consuming metabolic pathways or transporters, or to the presence of proton leakages from energized membranes. In addition, in natural environments, either terrestrial or aquatic, CO_2_ fixation is further limited by the concentration of atmospheric CO_2_, which can trigger photorespiration in C_3_ plants or induction of the carbon concentration mechanisms in C_4_ plants and algae. All these mechanisms would require additional energy for their activities (Osmond, 1981; Takabayashi *et al*., 2005; Walker *et al*., 2014; Burlacot *et al*., 2022), making alternative electron pathways of photosynthesis even more critical for sustaining plant and algal CO_2_ capture.

## Acknowledgements

The authors acknowledge the constructive comments on the manuscript from Dr. Joe Berry, Dr. Jennifer Johnson, Evan Saldivar, Leron Perez and technical support from Michel Philibert and Emmanuel Capra. This work was supported by the Carnegie Institution for Science (AB, AG), and the ANR grants OTOLHYD n°ANR-18-CE05-0029 (GP) and AlgalCCM n°ANR-22-CE44-0023-01 (GP, AM). This work was also in part supported by DOE award DE-SC0019417 to AG.

## Material and Methods

### Strains and growing conditions

*Chlamydomonas flvB* and *pgrl1* single mutants and their respective parental strains CC-4533 (for *flvB*), 137AH (for *pgrl1*) where previously described(Tolleter *et al*., 2011; Chaux *et al*., 2017). *flvB pgrl1*−1, −2 and −3 double mutant and their respective sibling strains WT2 and WT3 (used here as control strains) have been described previously(Burlacot *et al*., 2022) and are all from the progeny of a crossing of *flvB* and *pgrl1* performed in(Burlacot *et al*., 2022). Because *pgrl1 flvB* double mutants were obtained by crossing the two single mutants which have different genetic backgrounds (as described in (Burlacot *et al*., 2022)), the control strains used are siblings that resulted from the cross and that contained wild-type copies and levels of the FLVB and PGRL1 proteins. *pgrl1-2* is a mutant impaired in accumulation of the PGRL1 protein originating from crossing *flvB* and *pgrl1* performed in (Burlacot *et al*., 2022). All strains were grown photo-autotrophically in 125-mL flasks with cotton stoppers at 25 °C in a buffered minimal medium (20LmM MOPS pHL7.2) under constant illumination (80 µmol photons m^−2^ s^−1^) either under ambient CO_2_ or 2% CO_2_ in air. Amongst *pgrl1 flvB* double mutants and their control strains already described (Burlacot *et al*., 2022), *pgrl1 flvB*-1 and *pgrl1 flvB-*3 and WT2 and 3, respectively, were used for physiological measurements since their maximal photosynthesis was representative of other double mutants and control strains (Burlacot *et al*., 2022). All strains used in this study are available under the same names used here (except for *flvB* which is available as *flvB-21*) at the Chlamydomonas Resource Center (www.chlamycollection.org). Spectrum of the growing light used are shown in **Sup Fig. 13**.

### Gas exchange measurements

Gas exchange rates were measured using membrane inlet mass spectrometry (MIMS). Cells grown in air enriched or not at 2% CO_2_ were collected during the exponential phase by centrifugation at 450 g for 3 min and resuspended in 1.5 mL of fresh buffered minimal medium (pHL7.2) at 30 μg chlorophyll mL^-1^. HCO_3_^-^ was subsequently added to the cell suspension (10 mM final concentration). The cell suspension was then placed in the MIMS reaction vessel, 2 mL of ^18^O-enriched O_2_ (97% ^18^O Eurisop, France) were bubbled in the suspension. After closing the vessel, gas exchange was recorded during a dark to light transition (2,000 µmol photons m^−2^ s^−1^; green LEDs picked at 525nm; Luminus reference PT-121-G-L11-MPK). Light intensity was set to be saturating (**Sup Fig. 14**) using weakly absorbed green light for better homogeneity. Calculations of gross and net photosynthesis were done using the MIMS analysis software as previously described (Burlacot *et al*., 2020). In some experiments salicyl hydroxamic acid (SHAM, 400 µM final concentration) and/or myxothizol (Myxo, 2.5 µM final concentration) were added 1 min before the beginning of the measurements, except for the data presented in **Fig. 2A, B** where inhibitors were added during the measurement. Note here that SHAM has been reported to not inhibit the plastid terminal oxidase (Cournac *et al*., 2000). All Net O_2_, Net CO_2_, e^-^ quantum yield, e^-^ flow and Net photosynthesis sustained shown as histograms in **Figs. 2, 3, 4** and **Sup. Figs. 1, 7, 9, 11** are based on measurements after 9 minutes of illumination.

### Clover Expressing cells

137AH, WT2, *pgrl1*, *flvB*, *pgrl1flvB*-1 and −2 expressing the mitochondria targeted fluorophore Clover (mito-Clover) were generated as described in (Harmon *et al*., 2022). Briefly, cells were transformed with a linearized plasmid encoding mito-Clover and a hygromycin-resistance cassette using the Max Efficiency Reagent (Invitrogen). Resistant clones to hygromycin were screened for Clover fluorescence using a M1000 Tecan plate reader (Excitation: 488/7 nm; Emission: 515/10 nm) and mitochondrial signal was checked by confocal microscopy. Selected clones were grown at 80 µmol photons m^-2^ s^-1^ in 25 mL minimal medium in 125 mL flasks, in the presence of either normal or 2% CO_2_-enriched air until they reached a density of ∼10 µg chlorophyll mL^-1^. An aliquot of each culture was then briefly concentrated by centrifugation at 1,500g and cells layered on a 18-well microscope slide. Images were acquired using a TCS SP8 confocal laser-scanning microscope (Leica) and a ×63 objective. Acquisition settings were as follows: Clover excitation 488 nm, emission 500-547 nm; Chlorophyll excitation 488 nm, emission 694-733 nm. For homogeneity and clarity, signal intensities of each channel were adjusted using the Fiji software.

### Growth tests

The different *Chlamydomonas* strains were cultivated at air level of CO_2_ under moderate light (80 µmol photons m^-2^ s^-1^) at 25 °C. Cells were collected during exponential growth and diluted in fresh minimal medium to 0.125, 0.625 or 2.5L10^6^ cells mL^-1^. Seven-microlitre drops were spotted on 2% Agar plates of minimal medium (buffered with 20mM MOPS pHL7.2) with or without myxothiazol (2.5µM final concentration) and exposed to air enriched with 2% CO_2_. Homogeneous light was supplied by LED panels (see its spectrum in **Sup Fig. 13 B**), the different light intensities were obtained with various neutral filters. Temperature was maintained at 25L°C at the level of plates by means of fans.

### Electron quantum yield calculation

The e^-^ quantum yield of a given alternative electron pathway is defined as the proportion of electrons generated downstream PSI that are used for net photosynthesis when only this alternative electron pathway is active, following this equation:

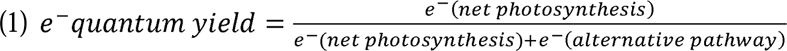

Where e^-^(net photosynthesis) is the flux of photosynthetic electrons directly used as reductants for net photosynthesis and e^-^(alternative pathway) is the flux of photosynthetic electrons through an alternative pathway. Because for PCEF and CMEF the sum of e^-^ (alternative pathway) and e^-^(net photosynthesis) equals gross photosynthesis, electron quantum yields of PCEF and CMEF were quantified using gross and net O_2_ evolution in mutants harbouring: only PCEF (*pgrl1* and *pgrl1-2* mutants treated with Myxo and SHAM), or only the AOX or the complex III/IV pathway of CMEF (*pgrl1 flvB*-1 and 3 mutants treated with Myxo or SHAM, respectively). The electron quantum yield was then calculated for each pathway using the following equation:

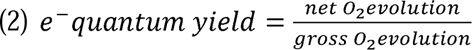

The theoretical electron quantum yield of CEF was calculated based on the assumption that CEF translocates 1.5 less protons per electron across the thylakoid membranes than PCEF (Kramer and Evans, 2011; Burlacot, 2023) and that this is the driving force leading to net photosynthesis energization. Note that this later assumption is well supported by the independent results from **Fig. 3 G** showing that the relative electron quantum yield of CMEF pathways and PCEF corresponds to their relative capacity at translocating protons across membranes. Hence, for the same net photosynthesis powered, CEF should be 1.5 times higher than PCEF, and the electron quantum yields of CEF could be written as:

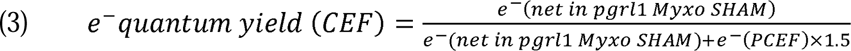

Where e^-^(net in *pgrl1* Myxo SHAM) is the net photosynthesis measured in the *pgrl1* mutant treated with Myxo and SHAM (hence having only PCEF active) and e^-^ (PCEF) being the O_2_ uptake rate in the same strain and condition.

Based on equation (1), we can rewrite e^-^(PCEF) as:

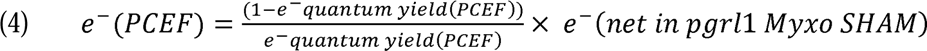

Hence, combining (3) and (4):

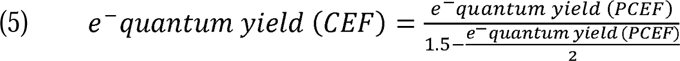

The electron quantum yield of CEF shown in **Fig. 3** is using equation (5) with the average electron quantum yield of PCEF. Note that the quantifications of the e^-^ quantum yield in *Chlamydomonas* assumes that removing two pathways (elimination of CEF and CMEF in the *pgrl1* mutant treated with inhibitors of respiration, and CEF and PCEF in the *pgrl1 flvB* double mutants) makes CO_2_ fixation dependent only on the third pathway. This dependency is supported by the finding that the *pgrl1 flvB* double mutant treated with inhibitors of respiration are unable to support any CO_2_ fixation, which also support the hypothesis that the contribution of other pathways, such as the NDA2-dependent CEF, is marginal.

### Calculation of electron flows via each pathway and their contribution to net photosynthesis in wildtype strains

The electron flow through PCEF and CMEF were deduced on wildtype strain using two independent methods based on measurements of gross O_2_ uptake made in **Sup. Fig. 2 and 3**. In the so called “mutant” method, CMEF was quantified as being the gross O_2_ uptake measured in the *flvB* mutant. Then, PCEF was quantified by subtracting the CMEF to the O_2_ uptake of CC-4533 (**Sup. Fig. S10**). This method was only used for CC-4533 since it is the only control strain for the *flvB* mutant. In the so called “inhibitor” method, PCEF was quantified as being the gross O_2_ uptake remaining in control strains treated with SHAM and Myxo. Then, CMEF was quantified by subtracting the PCEF to the O_2_ uptake in the same untreated control strain (**Sup. Fig. S10**). The calculated electron flows of PCEF and CMEF were then used for each strain to quantify the net photosynthesis powered by each of them following the equations derived from (1):

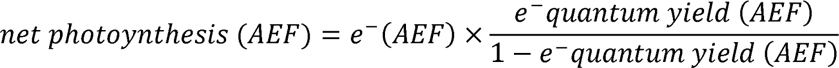

Where net photosynthesis (AEF) is the net photosynthesis sustained by a specific alternative electron flow, e^-^(AEF) being the electron flow through this AEF and e^-^ quantum yield (AEF) being the electron quantum yield of this alternative electron flow.

The total of net photosynthesis powered in a strain by PCEF and CMEF was then compared to the measured net photosynthesis in this same strain to quantify the “missing” net photosynthesis that had to be powered by CEF. Then the electron flow through CEF (e^-^(CEF)) was calculated by the following equation derived from (1):

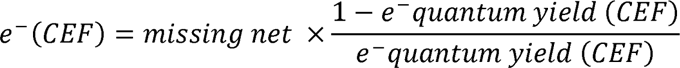

Where “missing net” is the calculated net photosynthesis that is not powered by PCEF and CMEF and e^-^ quantum yield (CEF) being the electron quantum yield of CEF.

### Protein extraction and Immunodetection

Cells were harvested by centrifugation at 1,000g for 5 minutes. Total cellular protein was extracted by heating the cell pellet at 100°C for one minute using an extraction aqueous solution composed of 1% SDS, 100 mM DTT, 1mM Aminocaproic acid, and 1mM benzamidine hydrochloride buffered with 60mM Tris (pH 6.8). The resulting cell lysate was then centrifuged at 10,000 g for 5 minutes to obtain the supernatant. The supernatant was subsequently incubated with acetone (80% final concentration) at −20°C for 1 hour. The protein pellet was collected by centrifugation at 5,000 g for 5 minutes and stored in aqueous solution containing 1% SDS, 5 mM DTT and buffered with 60mM Tris (pH 6.8). Protein content was estimated (Microplate BCA protein assay Kit-reducing agent compatible, Thermo Scientific) with bovine serum albumin as the standard. Protein extracts (10 µg protein) were loaded on a 12% SDS-PAGE gel and run for 1h at 150V in Tris-glycine SDS buffer and transferred onto a nitrocellulose membrane using semi-dry transfer (Transblot turbo, Biorad). Immunoblot detection was performed using antibodies raised against FlvB (1:1,000)(Chaux *et al*., 2017), and Pgrl1 (1:5,000)(Tolleter *et al*., 2011). Other antibodies against AOX1 (1:10,000) (AS06 152), COXIIB (1:10,000) (AS06 151) were obtained from Agrisera (https://www.agrisera.com/). An anti-rabbit HRP-conjugated antibody (1:50,000) (AS09 602) was used as a secondary antibody for immunodetection. Precision Plus Protein Dual Color Standards (Biorad) was used as the molecular weight marker. Uncropped and unprocessed scans of the blots shown in this study are visible in source data.

## Data Availability

Genes studied in this article can be found on https://phytozome-next.jgi.doe.gov/ under the loci Cre12.g531900 (FLVA), Cre16.g691800 (FLVB), Cre07.g340200 (PGRL1), Cre09.g395950 (AOX1), Cre03.g169550 (AOX2) and Cre01.g049500 (COXIIB).

