## Supplemental figures for "Green algae CO_2_ capture is powered by alternative electron pathways of photosynthesis"

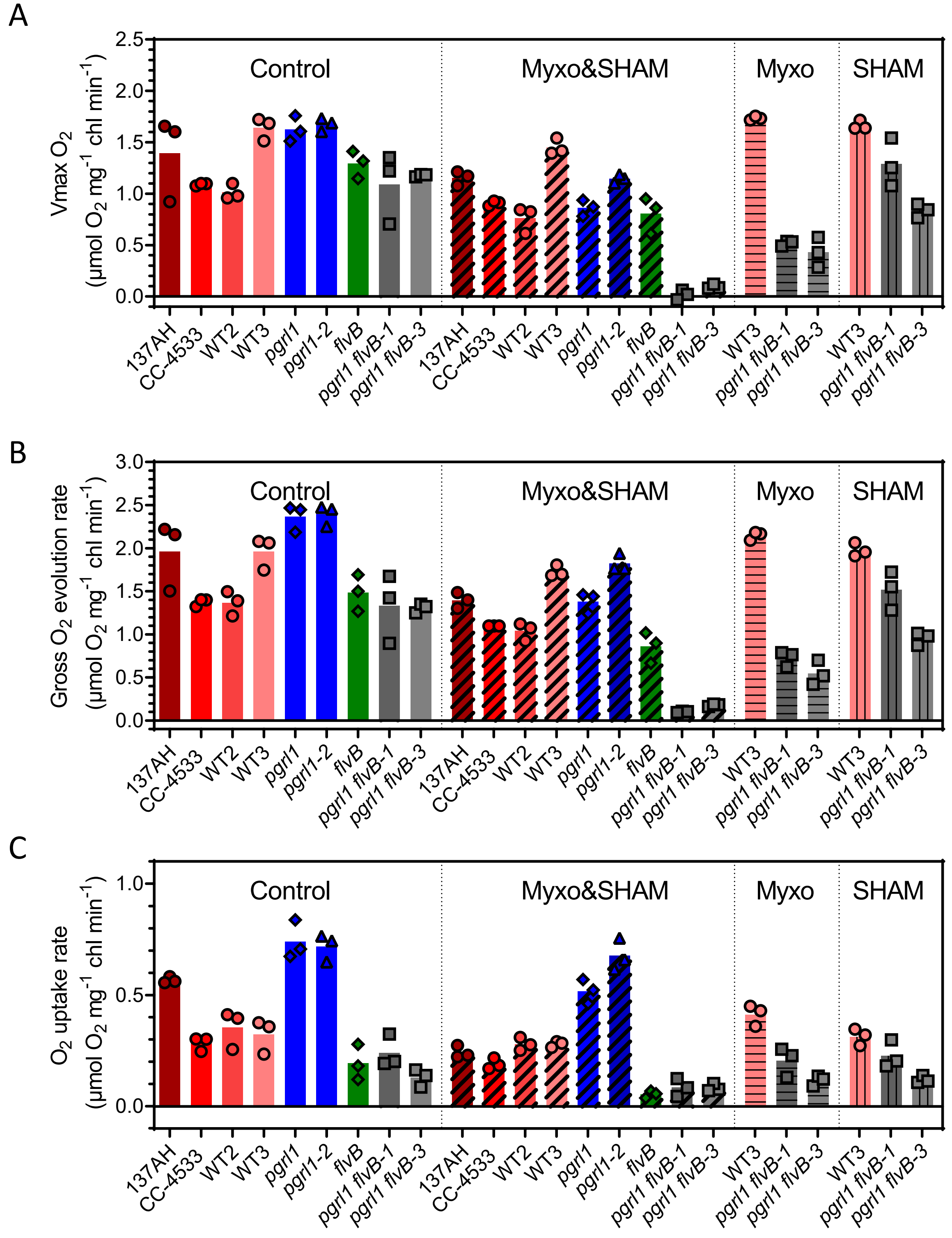


**Supplemental Figure S1. Effect of mitochondrial inhibitors on $O_2$ exchange rates on mutants and control strains grown at air level of $CO_2$.** $O_2$ exchange rates are measured from same experiments used in **Fig. 2 A-C**, **Sup Figs. S2, S3, S9, S11** and **S12**. Maximal net $O_2$ production (Vmax $O_2$) (**A**), gross $O_2$ evolution (**B**) and gross $O_2$ uptake (**C**) was measured on cells treated with myxothiazol (Myxo, 2.5 $\mu$M), salicylhydroxamic acid (SHAM, 400 $\mu$M), both Myxo and SHAM (Myxo&SHAM) or untreated (**Control**). Shown are the values measured after 9 minutes of illumination to reach steady state photosynthesis. Bars show the mean and dots show individual replicates (*n* = 3 biologically independent samples).

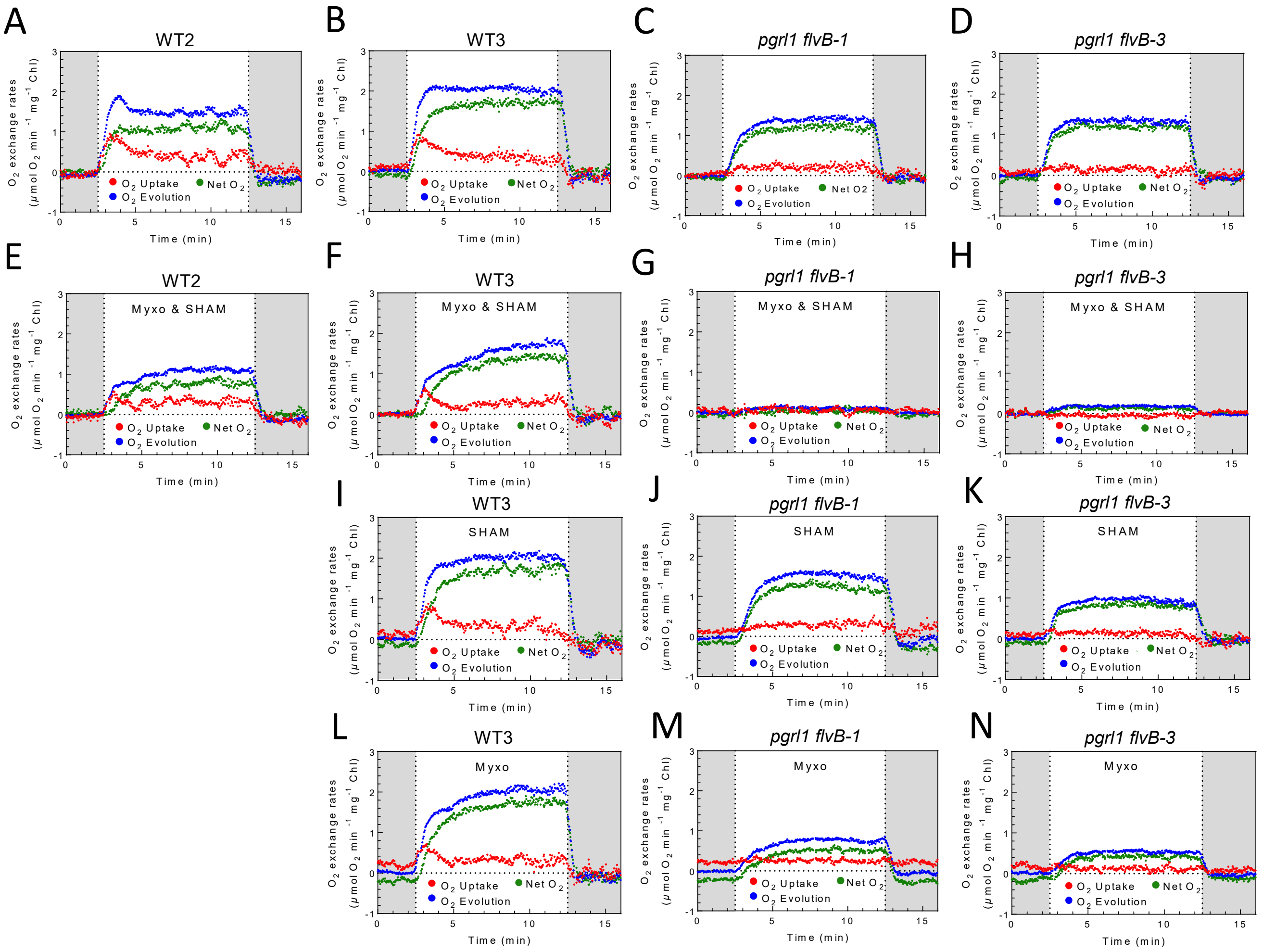


**Supplemental Figure S2. $O_2$ exchange rated in *pgrl1 flvB* double mutants and their control strains grown at air level of $CO_2$. A-N** Gross and net $O_2$ exchange rates upon a dark (grayed areas)-light-dark transition on cells treated with myxothiazol (Myxo, 2.5 $\mu$M) (**L**, **M**, **N**) , salicylhydroxamic acid (SHAM, 400 $\mu$M) (**I**, **J**, **K**) , both Myxo and SHAM (**E**, **F**, **G**, **H**) or untreated (**A**, **B**, **C**, **D**) . Cells were grown at air level of $CO_2$, shown are representative traces of n=3 biologically independent experiments for *pgrl1 flvB*-1 (**C**, **F**, **I**, **M**), *pgrl1 flvB-3* (**D**, **G**, **J**, **O**), *flvB* (**H**, **J**) and their control strains WT2 (**A**, **K**) and WT3 (**B**, **E**, **H**, **L**).

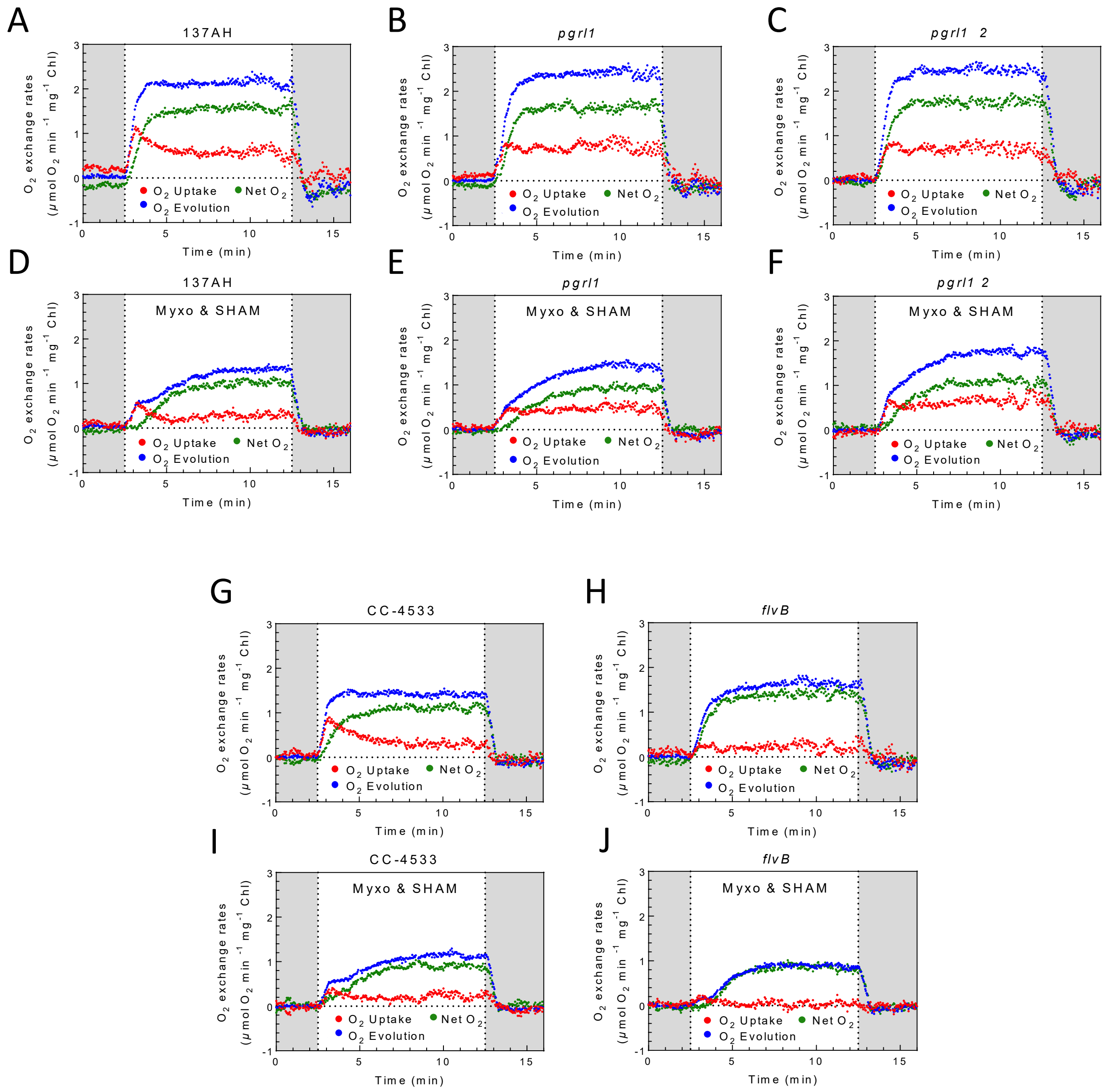


**Supplemental Figure S3. $O_2$ exchange rated in *pgrl1* and *flvB* single mutants and their control strains grown at air level of $CO_2$.** Gross and net $O_2$ exchange rates upon a dark (grayed areas) -light-dark transition on cells treated (**D**, **E**, **F**, **I**, **J**) or not (**A**, **B**, **C**, **G**, **H**) with myxothiazol (Myxo, 2.5 $\mu$M) and salicylhydroxamic acid (SHAM, 400 $\mu$M). Cells were grown at air level of $CO_2$, shown are representative traces of n=3 biologically independent experiments for *pgrl1* (**B**, **E**), *pgrl1* 2 (**C**, **F**), *flvB* (**H**, **J**) and their respective control strains 137AH (**A**, **D**) and CC-4533 (**G**, **I**).

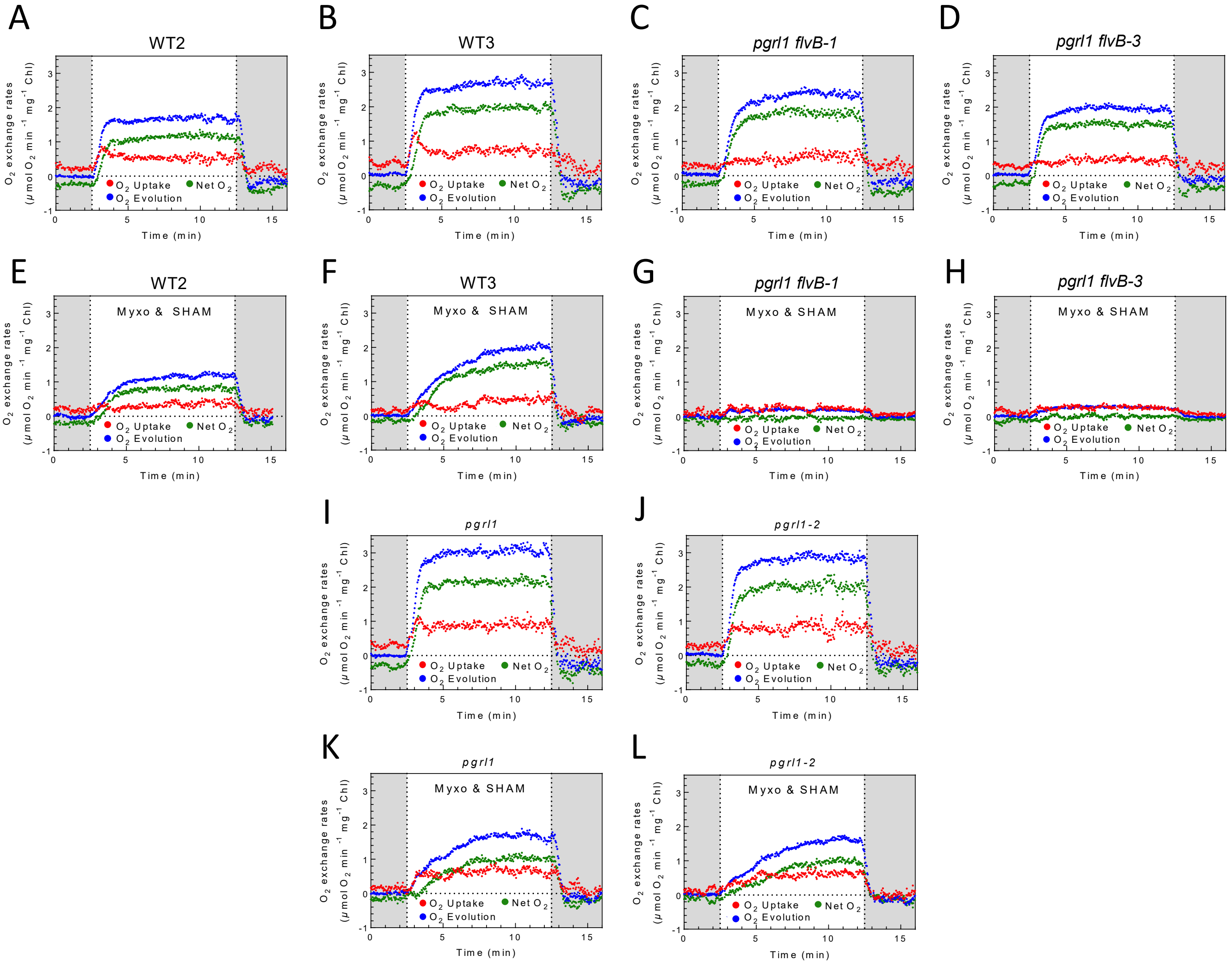


**Supplemental Figure S4. $O_2$ exchange rated in *pgrl1, pgrl1 flvB* and their control strains grown in air enriched with 2% $CO_2$.** Gross and net $O_2$ exchange rates upon a dark (grayed areas) -light-dark transition on cells treated (**E**, **F**, **G**, **H, K, L**) or not (**A**, **B**, **C**, **D**, **K**, **L**) with myxothiazol (Myxo, 2.5 $\mu$M) and salicylhydroxamic acid (SHAM, 400 $\mu$M). Cells were grown with air enriched with 2% $CO_2$, shown are representative traces of n=3 biologically independent experiments for *pgrl1 flvB*-1 (**C**, **G**), *pgrl1 flvB-3* (**D**, **H**), their respective control strains WT2 (**A**, **E**) and (**B**, **F**), *pgrl1* (**I**, **K**) and *pgrl1* 2 (**J**, **L**).

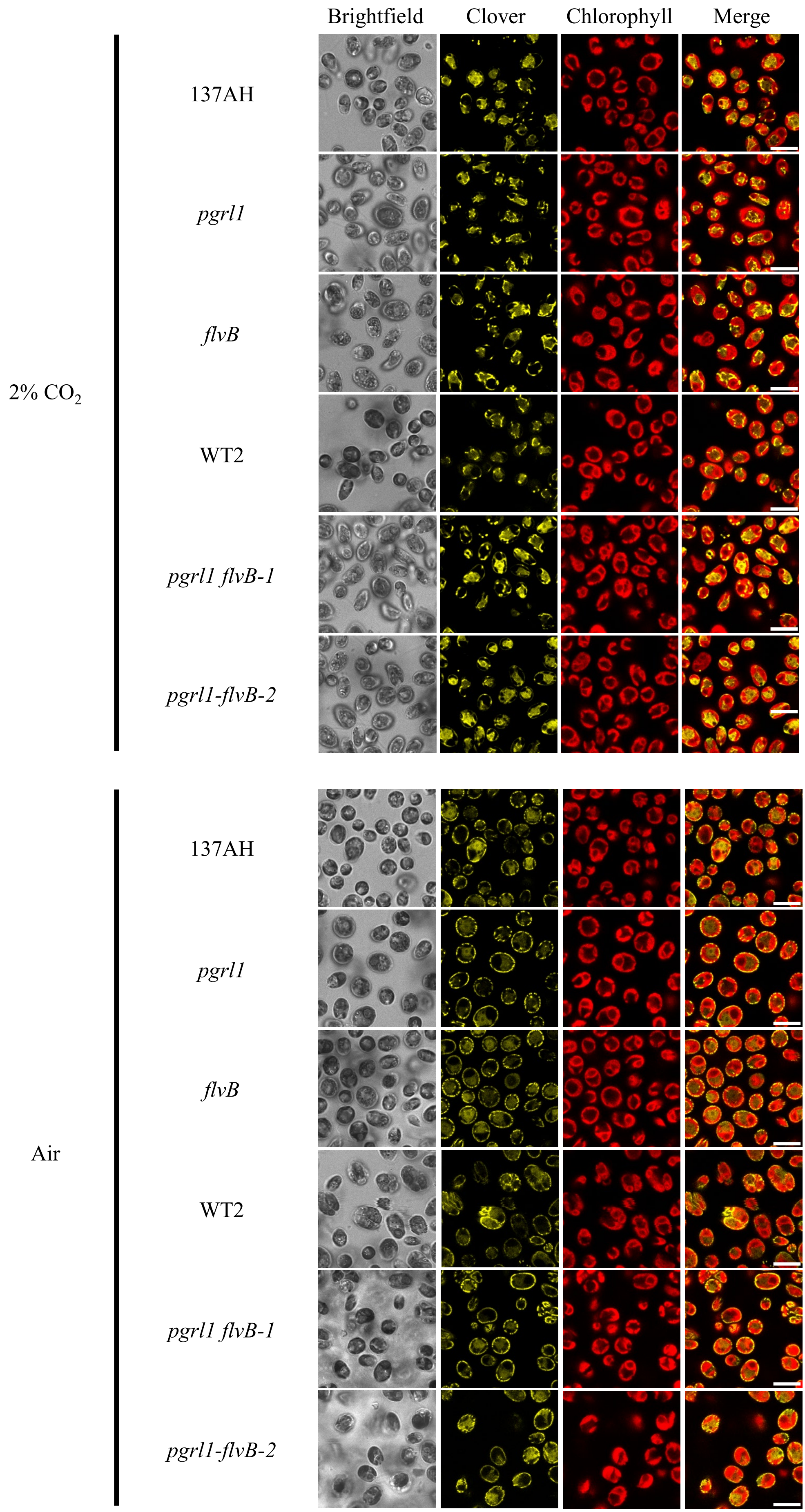


**Supplemental Figure S5. Localization of mitochondria in *pgrl1*, *flvB*, *pgrl1 flvB-1* and -2 and their control strain in air enriched or not with 2% $CO_2$.** Cross section images showing subcellular localization of the mitochondria-localized mito-Clover and chlorophyll fluorescence in *pgrl1*, *flvB*, *pgrl1 flvB-1, pgrl1 flvB-2* and the control strains 137AH and WT2, grown in air or 2% $CO_2$-enriched air. Shown are representative of two independent experiments.

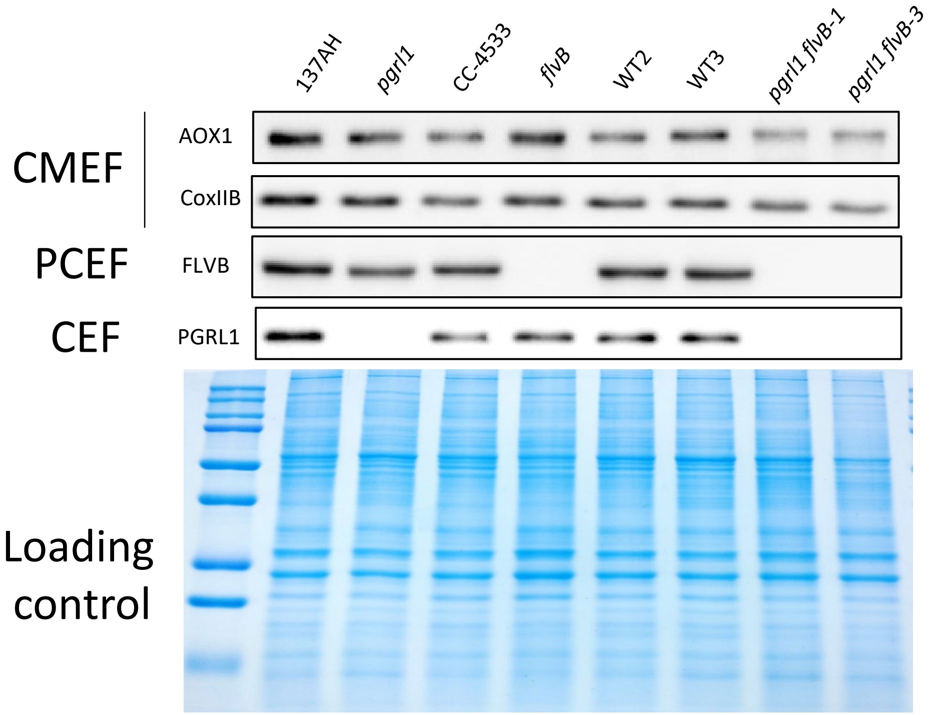


**Supplemental Figure S6. Immunodetection of proteins involved in CMEF, PCEF and CEF in the strains used in this study.** Immunodetection of PGRL1, FLVB, AOX1, CoxIIB in pgrl1, flvB mutants and *pgrl1 flvB* double mutants and their respective controls grown at air level of $CO_2$.

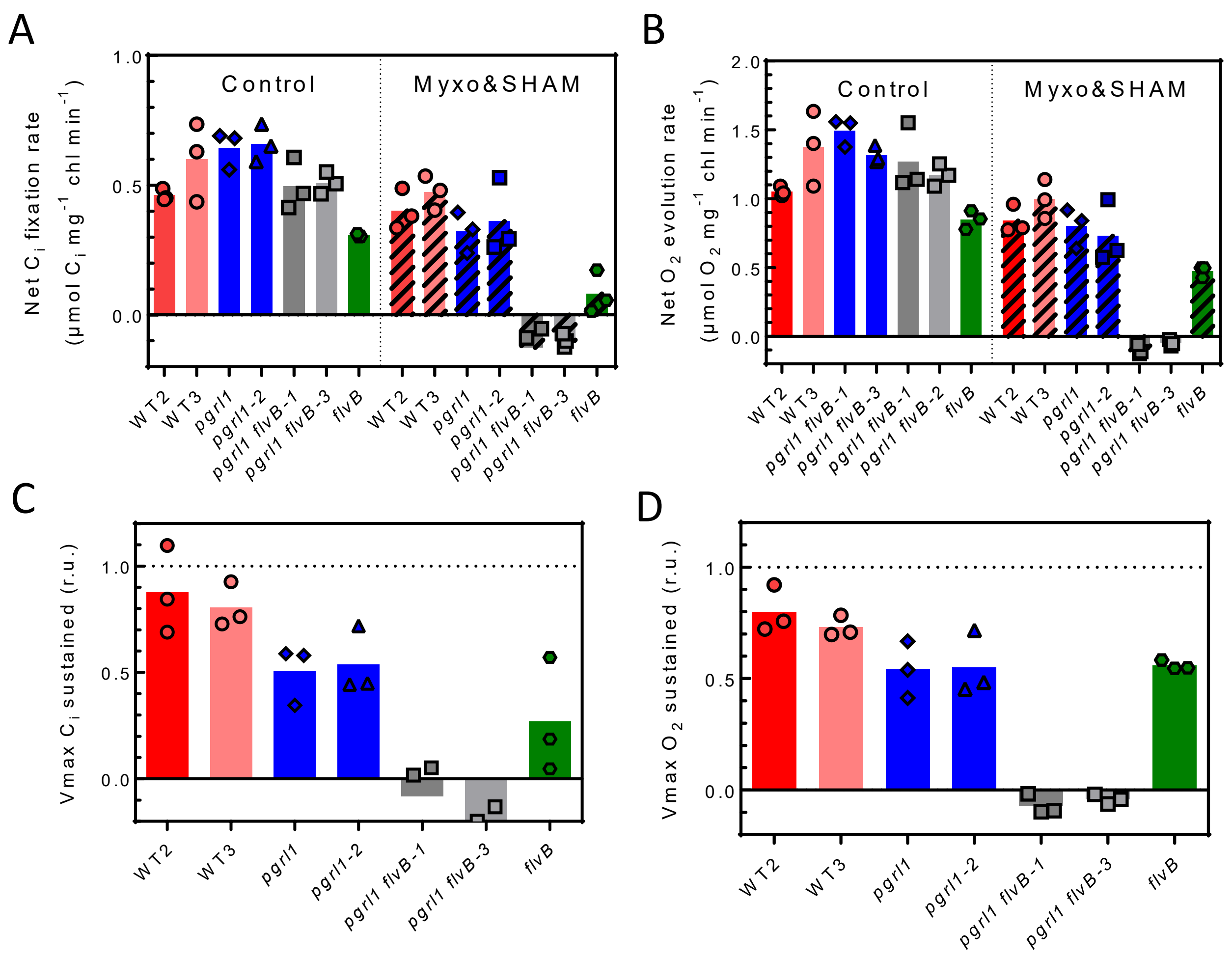


**Supplemental Figure S7. Comparison of net inorganic carbon fixation and net $O_2$ production.** Inorganic carbon ($C_i$) and $O_2$ exchange rates are measured concomitantly during a dark to light transition. Data shown are from same experiments used in **Sup Figs. S8**. **A**, **B** Net $O_2$ evolution (**B**) and $C_i$ fixation (**A**) on cells treated (right panels) or not (left panels) with myxothiazol (Myxo, 2.5 $\mu$M) and salicylhydroxamic acid (SHAM, 400 $\mu$M). **C, D** Maximal net $C_i$ fixation (**C**) and net $O_2$ evolution (**D**) sustained after treatment with myxothiazol and salicylhydroxamic acid. Cells were grown at air level of $CO_2$, shown are representative traces of n=3 biologically independent experiments for *pgrl1*, *pgrl1* 2, *flvB, pgrl1 flvB-1, pgrl1 flvB-3* and the control strains WT2 and WT3.

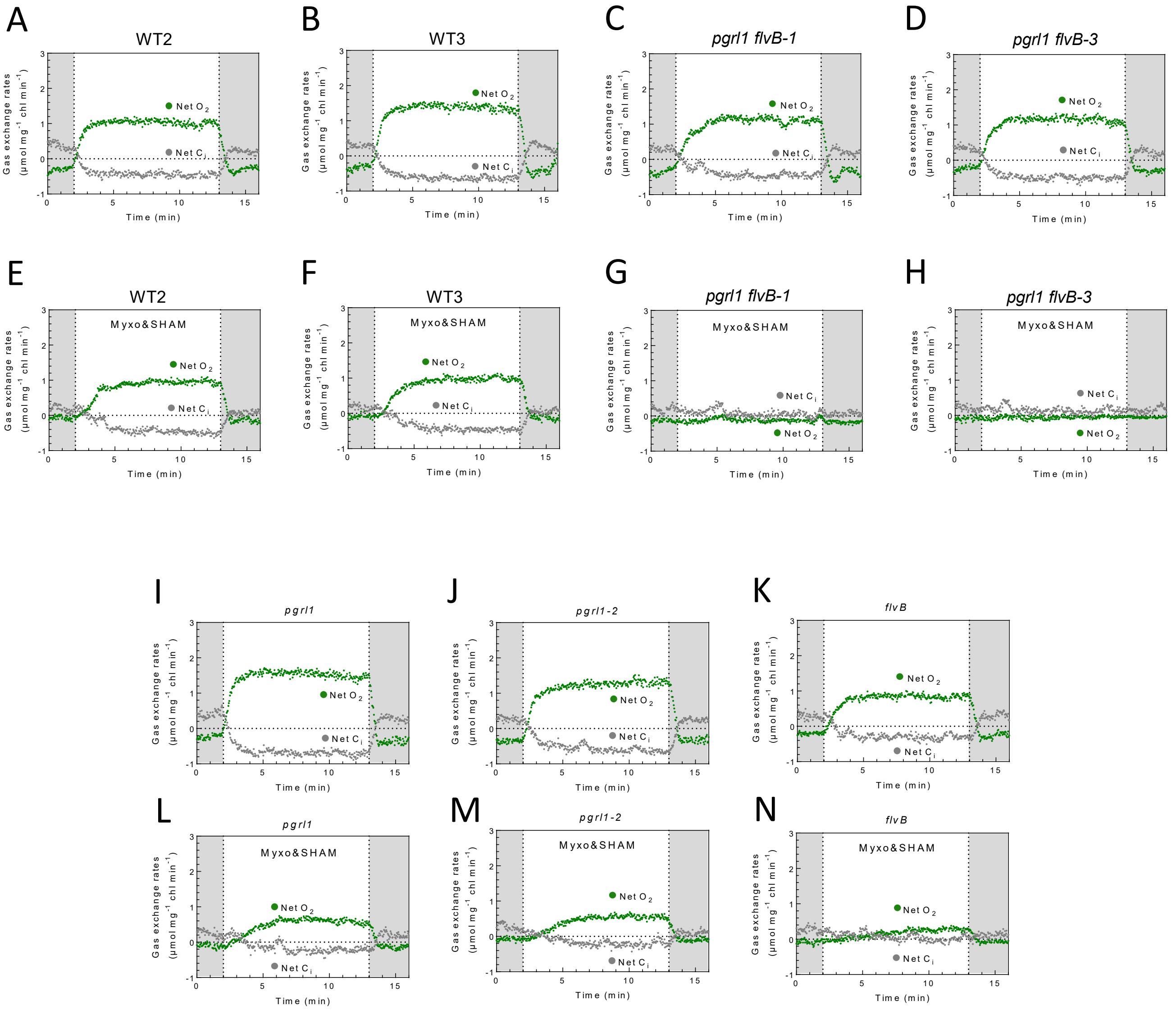


**Supplemental Figure S8. Net $C_i$ and $O_2$ exchange rated in *pgrl1* and *flvB* single and double mutants and the control strains WT2 and WT3.** Inorganic carbon ($C_i$) and $O_2$ exchange rates are measured concomitantly upon a dark (grayed areas) -light-dark transition on cells treated (**E**, **F**, **G**, **H, L, M, N**) or not (**A**, **B**, **C**, **D**, **I**, **J**, **K**) with myxothiazol (Myxo, 2.5 $\mu$M) and salicylhydroxamic acid (SHAM, 400 $\mu$M). Cells were grown at air level of $CO_2$, shown are representative traces of n=3 biologically independent experiments for *pgrl1* (**I**, **L**), *pgrl1* 2 (**J**, **M**), *flvB* (**K**, **N**), *pgrl1 flvB-1* (**C**, **G**), *pgrl1 flvB-3* (**D**, **H**) and the control strains WT2 (**A**, **E**) and WT3 (**B**, **F**).

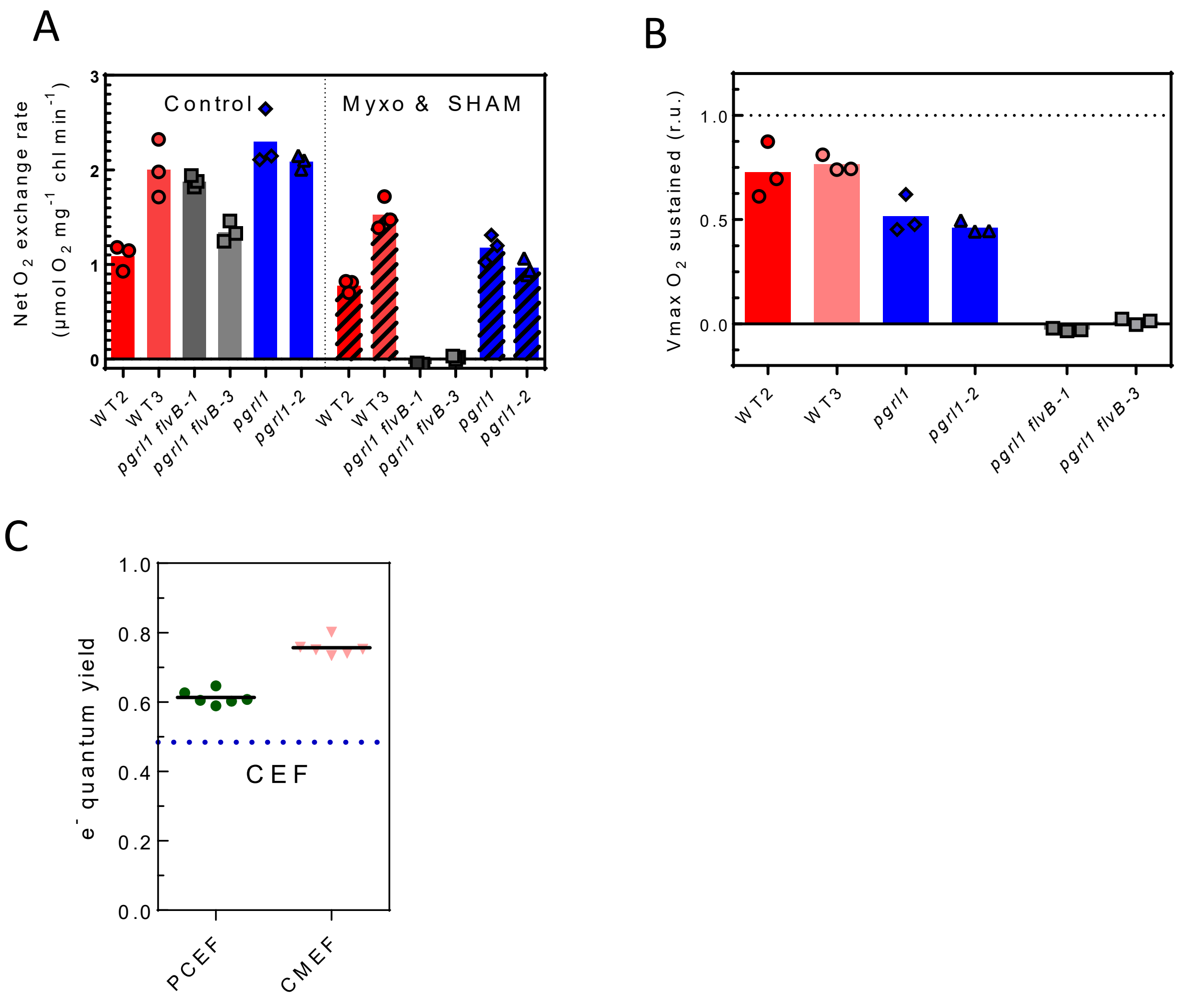


**Supplemental Figure S9. Net $O_2$ exchange rates characteristic and $e^-$ quantum yield measured in strains grown in air enriched with 2% $CO_2$.** Data shown are obtained from the same experiments used in **Fig. 2 D**, **Sup Fig. S4**. **A**, Net $O_2$ evolution on cells treated (right panels) or not (left panels) with myxothiazol (Myxo, 2.5 $\mu$M) and salicylhydroxamic acid (SHAM, 400 $\mu$M). **B** Maximal net $O_2$ evolution sustained after treatment with myxothiazol and salicylhydroxamic acid. **C** Electron quantum yield of PCEF (green), CMEF (pink) and CEF (dotted line). Cells were grown in air enriched with 2% $CO_2$, Bars show the mean and dots show individual replicates (*n* = 3 biologically independent samples).

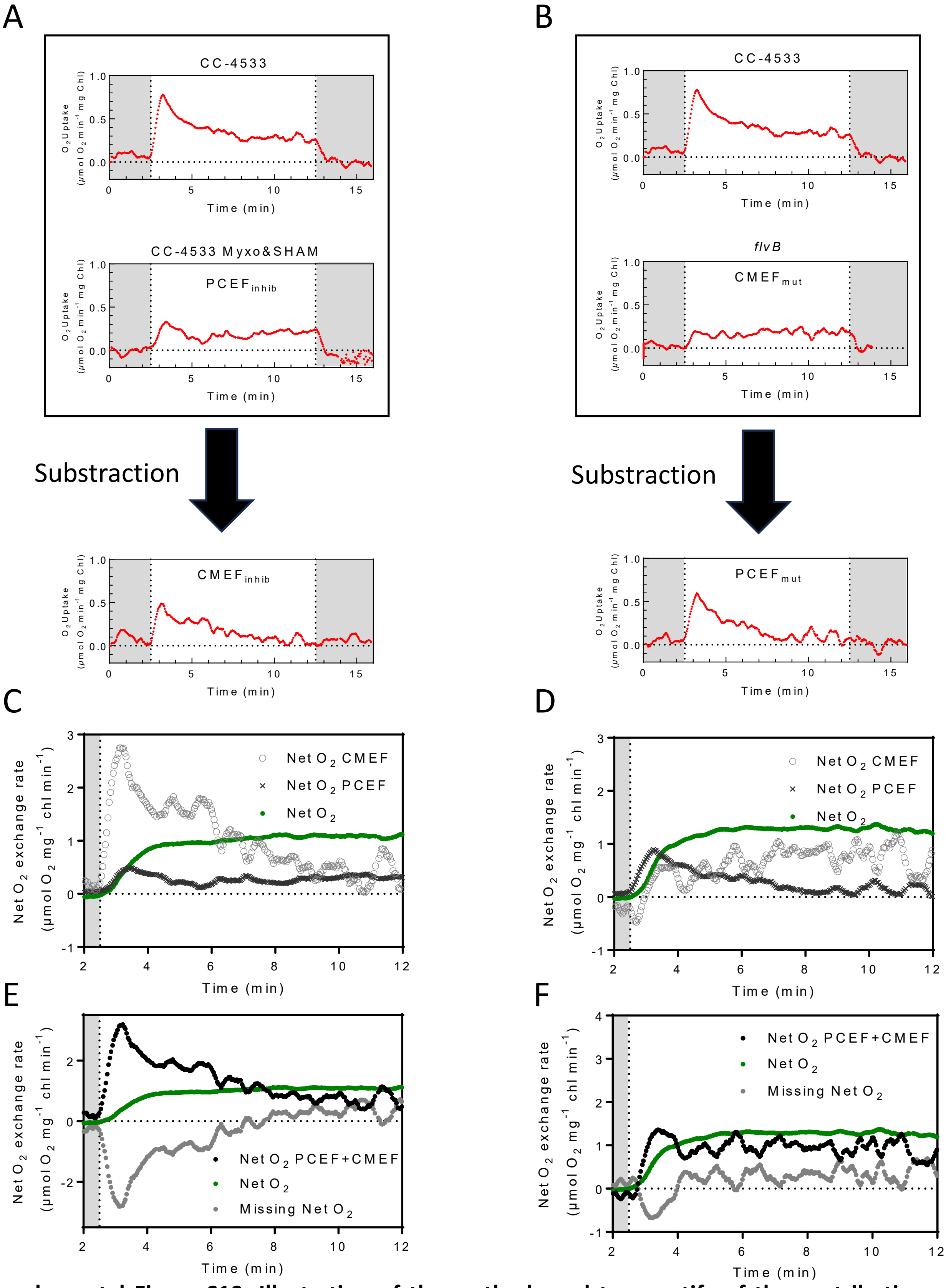


**Supplemental Figure S10. Illustration of the method used to quantify of the contribution of each photosynthetic electron flow to $CO_2$ fixation.** Illustrated here are the two different methods used, one based on the use of inhibitors (inhib) (**A**, **C**, **E**) the other on mutants (mut) (**B**, **D**, **F**). The first step for both methods is to calculate the gross $O_2$ uptake mediated by CMEF and PCEF either by first quantifying $PCEF_{inhib}$ and deducting $CMEF_{inhib}$ (**A**), or by first quantifying $CMEF_{mut}$ and deducting $CPEF_{mut}$ (**B**). Then, those fluxes are combined with the $e^-$ quantum yield of each flux to predict the net $O_2$ evolution powered by CMEF (gray cirlces) and PCEF (gray crosses) (**C**, **D**). The net $O_2$ hence powered by CMEF and PCEF (gray dots) is then compared to the actual net $O_2$ evolution measured (green dots) to compute the net $O_2$ evolution that should be powered by CEF (missing Net $O_2$) (**E**, **F**). Shown here are average signals of $n$ = 3 biologically independent samples.

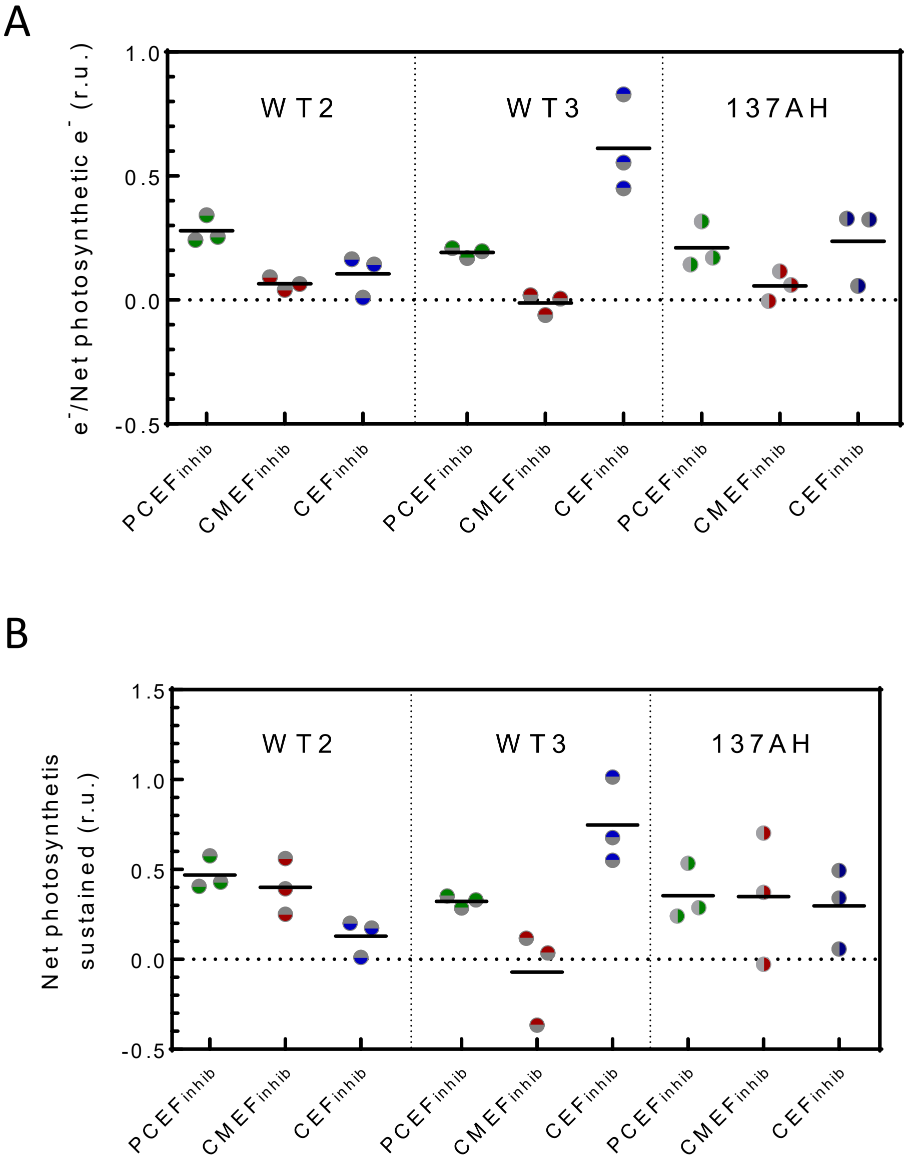


**Supplementatl Figure S11. Quantification of the contribution of each photosynthetic electron flow to net photosynthesis.** Data used for this calculation are the same as the one in **Supplemental Figs. S1, S2, S3.** Evaluation of the electron flow through PCEF, CMEF and CEF relative to LEF (**A**), and net photosynthesis sustained by PCEF, CMEF and CEF (**B**) based on the inhibitor method (**Sup. Fig. 12**) on WT2, WT3 and 137AH. Because of an twice greater dark $O_2$ uptake in 137AH, CMEF of 137AH was lowered by the dark $O_2$ uptake. Cells were grown at air levels of $CO_2$, Bars show the mean and dots show individual replicates (*n* = 3 biologically independent samples).

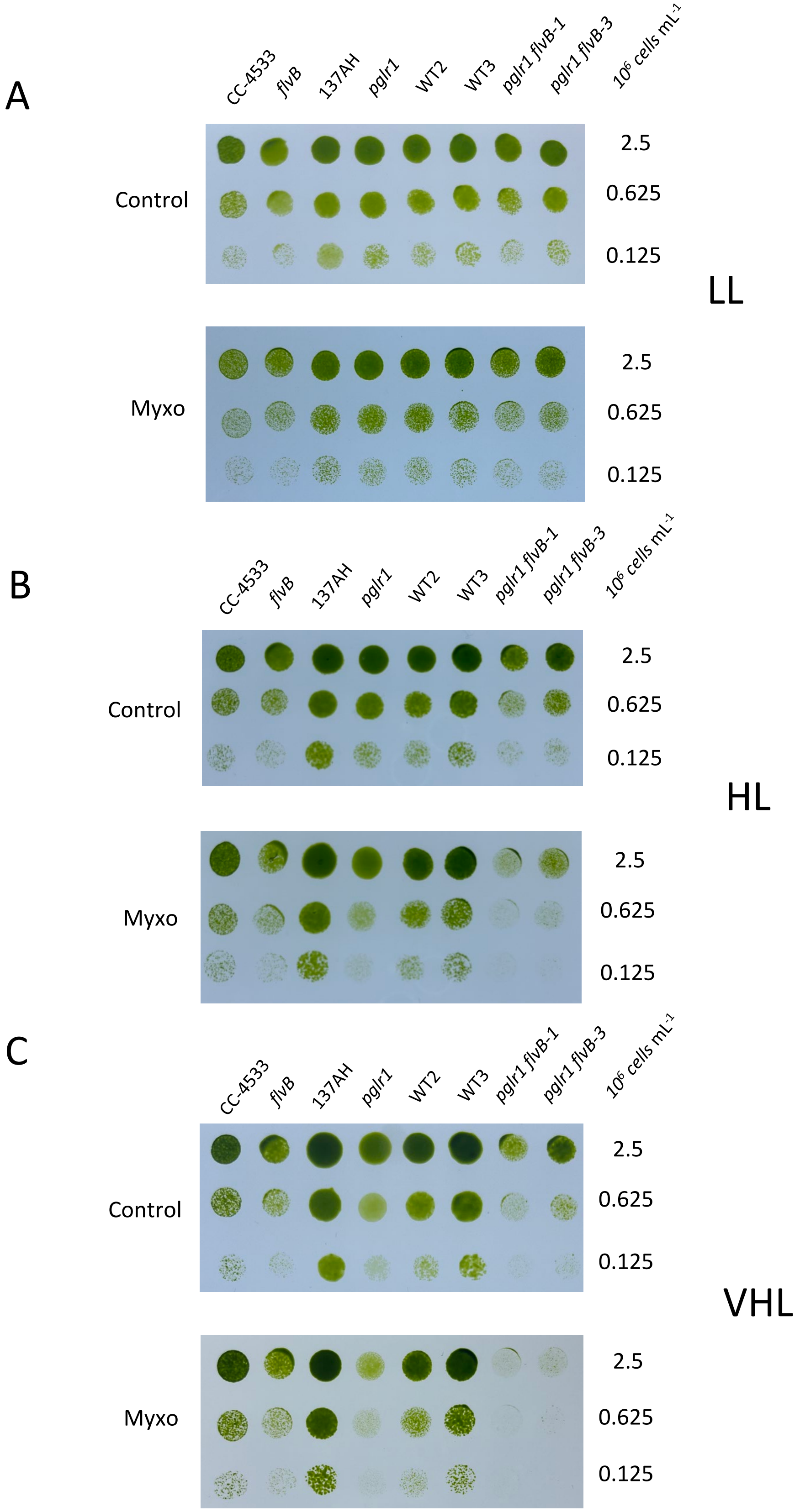


**Supplemental Figure S12. Growth of mutant and control strains at various light intensities.** Cells were spotted on plates containing minimal medium in the absence (control) or presence (Myxo) of myxothiazol (2.5µM) and grown under low light (LL, 30 µmol photons $m^{-2}$ $s^{-1}$), high light (HL, 100 µmol photons $m^{-2}$ $s^{-1}$) or very high light (VHL, 200 µmol photons $m^{-2}$ $s^{-1}$). Cells were grown in air enriched with 2% of $CO_2$ at 25°C. Growth was assessed after three days in *pgrl1*, *flvB*, and their respective control strains (137AH and CC-4533) and on double mutants (*pgrl1 flvB*-1 -3) and their control strains (WT2, WT3). Shown are representative spot tests of n = 10 independent experiments.

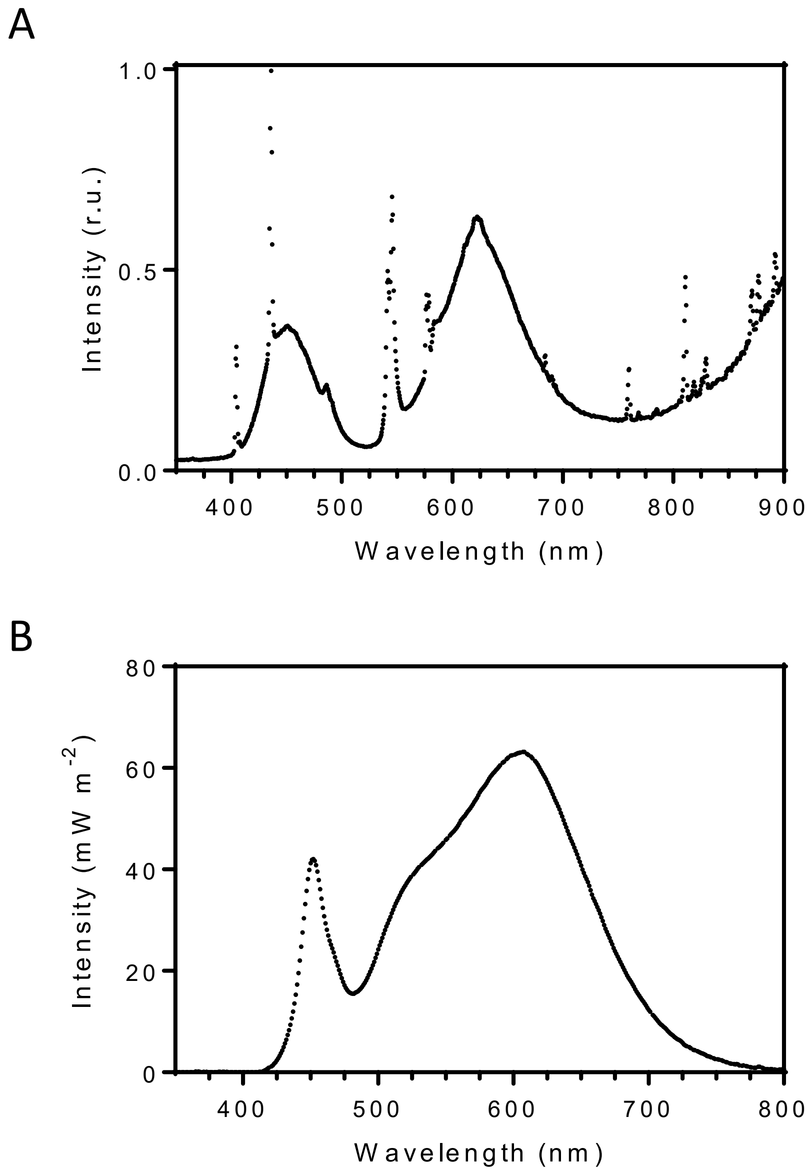


**Supplemental Figure S14. Growing light spectra.** Light spectra of light sources used for liquid cultures throughout the manuscript (**A**) or for cultures on solid media (**B**) in **Fig. 3 C** and **Sup Fig. 12**.

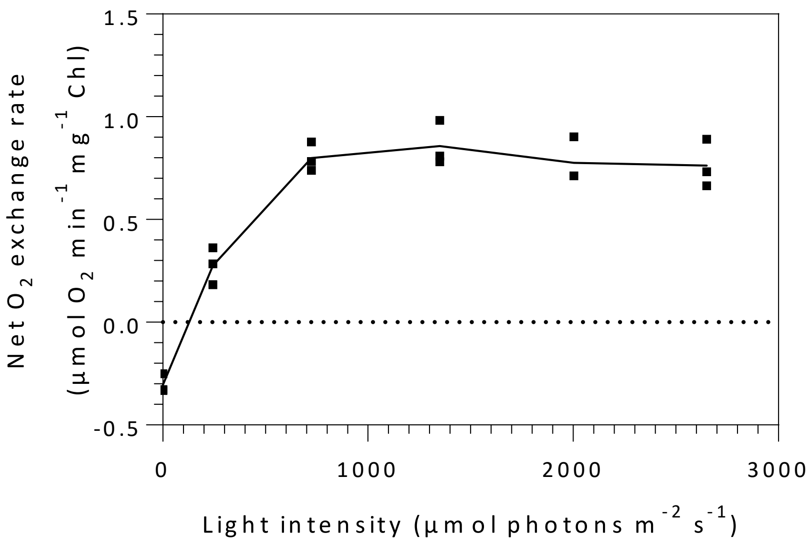


**Supplemental Figure S15. Light saturation curve of net $O_2$ exchange rate measured with green actinic light.** Cells were grown at air $CO_2$ level, collected during the exponential phase by centrifugation and resuspended in 1.5 mL of fresh buffered minimal medium (pH 7.2) at 30 µg chlorophyll $mL^{-1}$. $NaHCO_3^-$ (10 mM final concentration) was added to the cell suspension. After 2 minutes dark acclimation in the MIMS cuvette, green light was switched on and gradually increased every 3 minutes. Connecting line shows the mean and dots show individual replicates (*n* = 3 biologically independent samples)

### Source Data

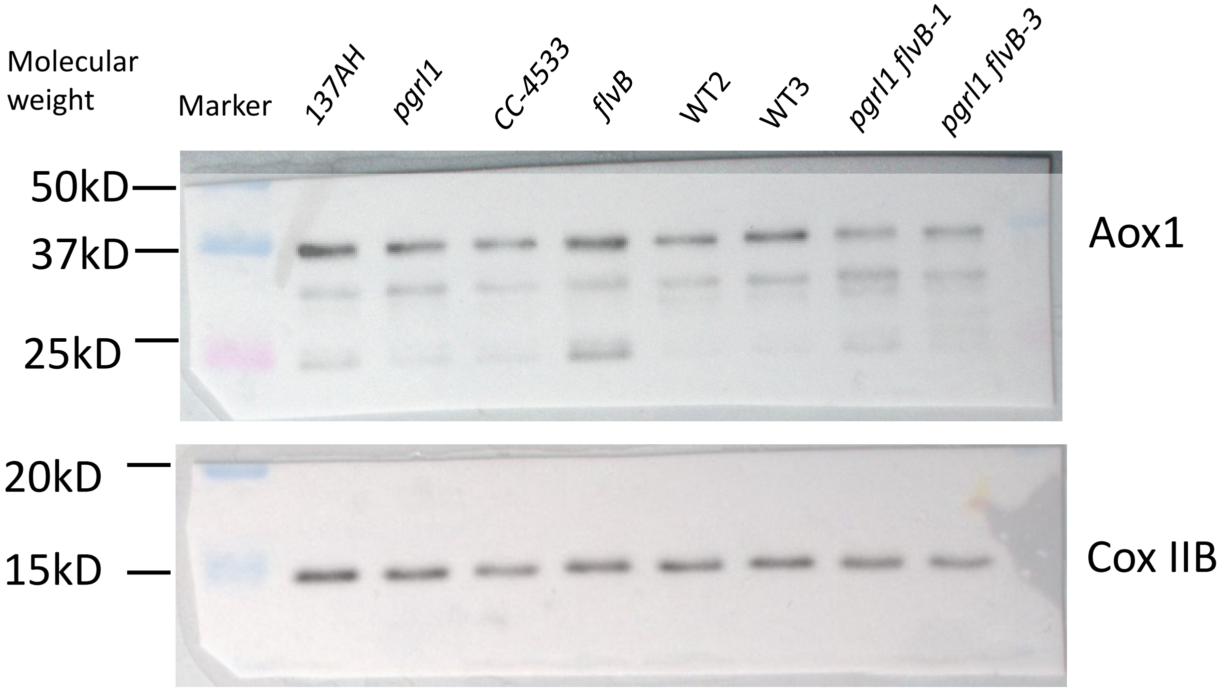
Molecular weight
Marker
137AH
pgrl1
CC-4533
flvB
WT2
WT3
pgrl1 flvB-1
pgrl1 flvB-3
50kD
37kD
25kD
Aox1
20kD
15kD
Cox IIB

Aox 1

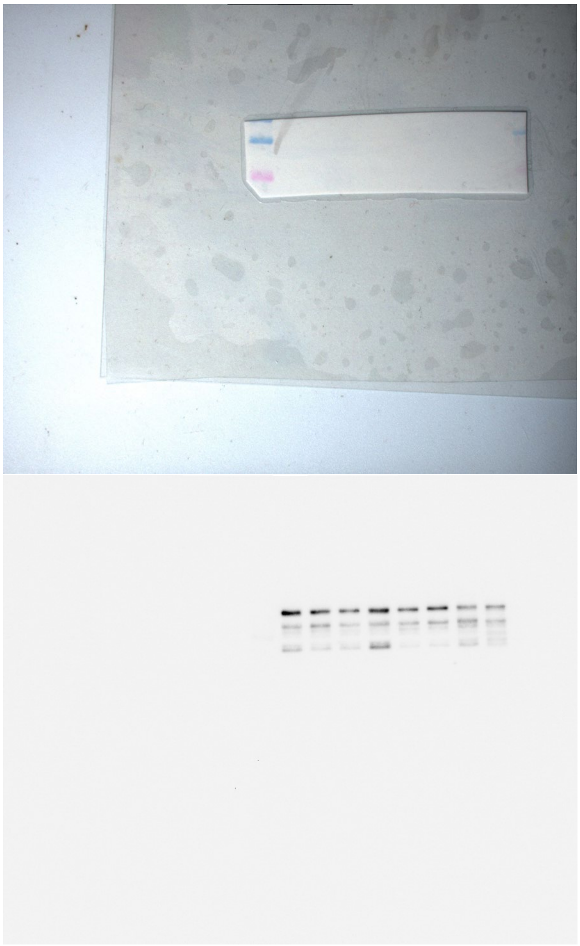

Cox IIB

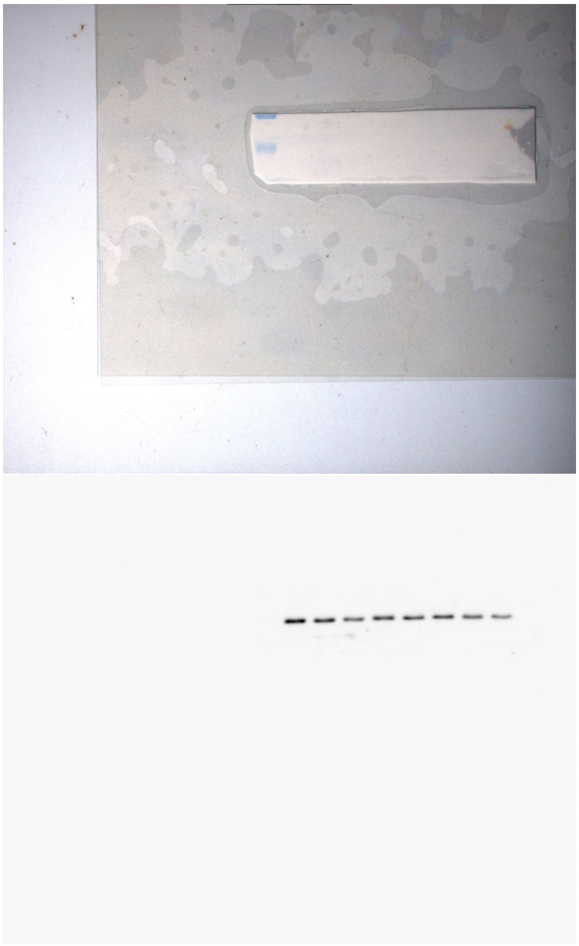

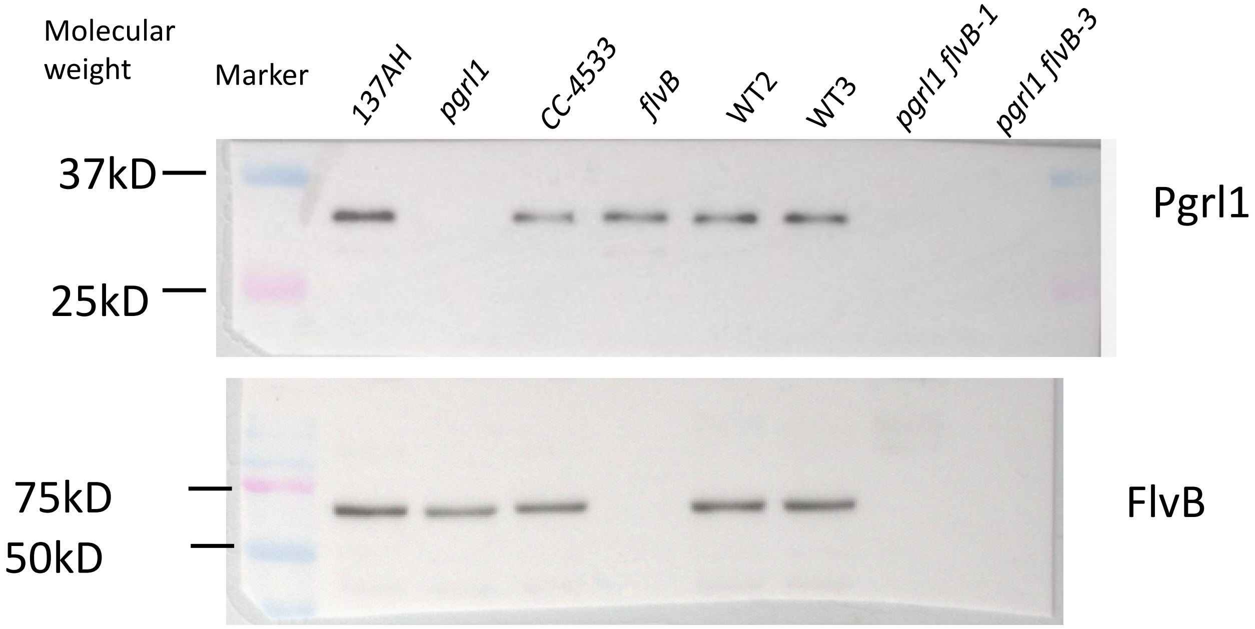
Molecular weight
Marker
137AH
pgrl1
CC-4533
flvB
WT2
WT3
pgrl1 flvB-1
pgrl1 flvB-3
37kD
25kD
Pgrl1
75kD
50kD
FlvB

FlvB

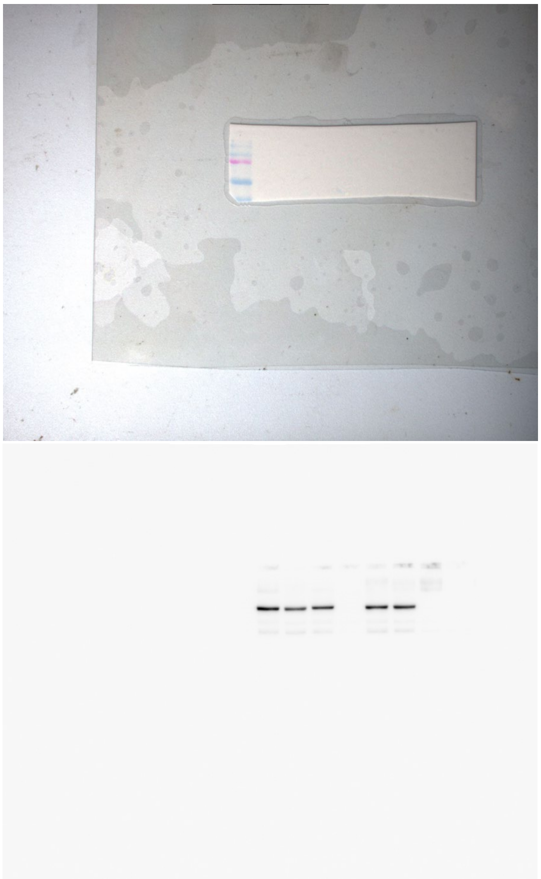

Pgrl1

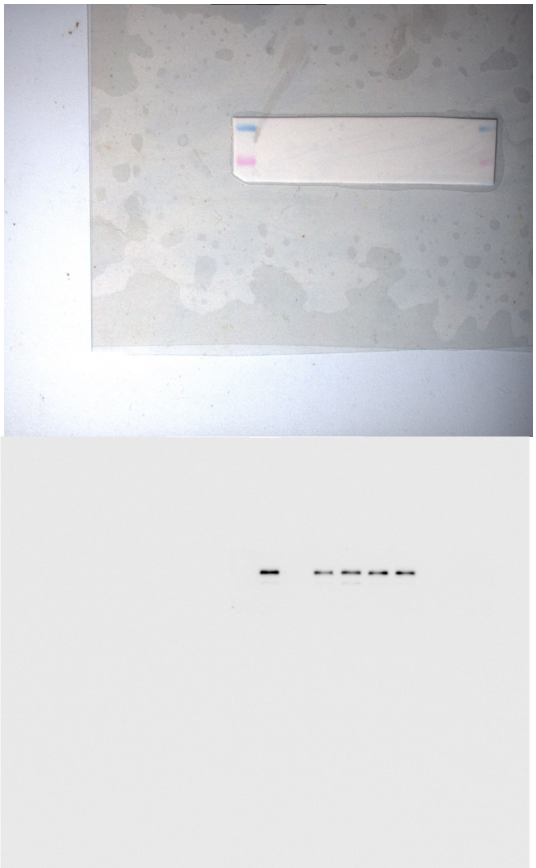
